# Sex-dependent gating of astrocyte-driven nociceptive plasticity

**DOI:** 10.64898/2026.11.18.767473

**Authors:** Laura Klinger, Lukas Uhrik, Eva Ravagnan, Raphael Holzinger, Lidia Trofimova, Danijela Kurija, Anna SM Siegert, Sibel Ada, Petra Waidhofer-Söllner, Manuela Schmidt, Ruth Drdla-Schutting

## Abstract

Sex differences in pain are well established, yet how nociceptive processing is differentially regulated across sexes remains poorly understood. Here, we show that spinal astrocytes provide pathway-specific control of acute nociceptive synaptic plasticity in rats. Chemogenetic activation of distinct astrocytic G-protein-coupled receptor pathways induced synaptic depression or potentiation at spinal nociceptive synapses in males, but failed to evoke detectable plasticity in females. Despite this lack of astrocyte-induced synaptic plasticity, females developed mechanical hypersensitivity comparable to males, revealing a dissociation between synaptic plasticity and pain-related behavior. Gonadectomy prevented astrocyte-induced plasticity in males, whereas testosterone treatment enabled Gq-dependent synaptic depression in ovariectomized females. Females exhibited more prominent perineuronal net labeling, and enzymatic degradation of extracellular matrix (ECM) components enabled Gq-dependent synaptic depression in females. Together, these findings identify hormonal status and ECM organization as important determinants of the susceptibility of spinal nociceptive circuits to astrocyte-driven synaptic plasticity.

## INTRODUCTION

Spinal nociceptive processing arises from dynamic interactions between neuronal and non-neuronal elements within the spinal dorsal horn (SDH) and across broader pain pathways. While nociceptive transmission and plasticity in the SDH have been extensively characterized at the synaptic and circuit level, mechanistic insight has largely emerged from a neuron-centric perspective. By contrast, glial contributions to pain have primarily been studied in chronic and pathological conditions. Most current understanding of glial function derives from neuropathic or inflammatory models, in which astrocytes and other immune-competent cells contribute to the development and maintenance of persistent pain through sustained signaling and neuroinflammatory processes (*1*). However, how acute glial modulation of spinal nociceptive circuits varies between sexes, and which factors determine the resulting synaptic changes under otherwise basal conditions, remains poorly understood (*2*).

Astrocytes are well positioned to contribute to sex-dependent differences in nociceptive processing. Sex differences in astrocyte number, morphology, gene expression, and functional properties have been reported across multiple regions of the central nervous system (*3–5*), while astrocytes are increasingly recognized as active regulators of synaptic transmission and plasticity (*6–8*). Through interactions with synapses, regulation of extracellular ion and transmitter homeostasis, intracellular calcium signaling, and the release of neuromodulatory factors, astrocytes can influence the functional state of surrounding neuronal networks (*9*). Importantly, the consequences of astrocyte activation may depend not only on signaling within astrocytes themselves, but also on the neuronal and extracellular environment within which these signals act (*10*). Such context-dependent constraints may provide one explanation for why similar astrocytic activation produces different effects across biological contexts, including between sexes.

We recently described a form of astrocyte-driven synaptic plasticity, termed astrocytogenic synaptic depression (astroSD) (*11*). Strikingly, this form of astrocyte-driven plasticity was observed in male but not female rats. This finding raised the question of whether the absence of synaptic plasticity in females reflects sex differences in astrocyte responsiveness itself or a sex-dependent restriction in the translation of astrocyte activity into neuronal plasticity.

Here, we investigated the conditions under which acute astrocyte activation engages plasticity in spinal nociceptive circuits. Using selective activation of astrocytic Gq-and Gi-coupled signaling pathways, we demonstrate that astrocytes induce opposing forms of synaptic plasticity in males, whereas neither form is detectable in females under basal conditions, despite astrocytic calcium responses in both sexes. Moreover, astrocyte activation induces mechanical hypersensitivity in both sexes, revealing a dissociation between behavioral sensitivity and astrocyte-driven synaptic plasticity. We further identify hormonal state and ECM organization as factors regulating the susceptibility of nociceptive synapses to astrocyte-driven plasticity. Together, our findings reveal sex-dependent gating of acute astrocyte-induced synaptic plasticity in the SDH and indicate that the consequences of astrocytic signaling depend on the physiological and extracellular context of nociceptive circuits.

## RESULTS

### Astrocyte-driven synaptic plasticity is bidirectional and sex-dependent

To selectively engage astrocytic GPCR signaling in the SDH, we used AAV-mediated expression of astrocyte-targeted DREADDs and recorded C-fiber-evoked field potentials in the superficial laminae after sciatic nerve stimulation in deeply anaesthetized rats (**Fig. 1a**). DREADD expression showed comparable distribution and astrocytic selectivity across constructs and sexes (**Extended Data Fig. 1**). In males, selective activation of astrocytic Gq-DREADDs by i.v. clozapine N-oxide (CNO, 3 mgꞏkg^-1^) induced robust and long-lasting depression of synaptic strength, measured as field potential area in the SDH (i.e. astroSD) ((*11*); **Fig. 1b**). In females, the same manipulation failed to induce detectable plasticity ((*11*); **Fig. 1c**), and CNO had no effect in animals transduced with a sham virus lacking DREADD expression ((*11*); **Extended Data Fig. 2a**).

**Fig. 1.**
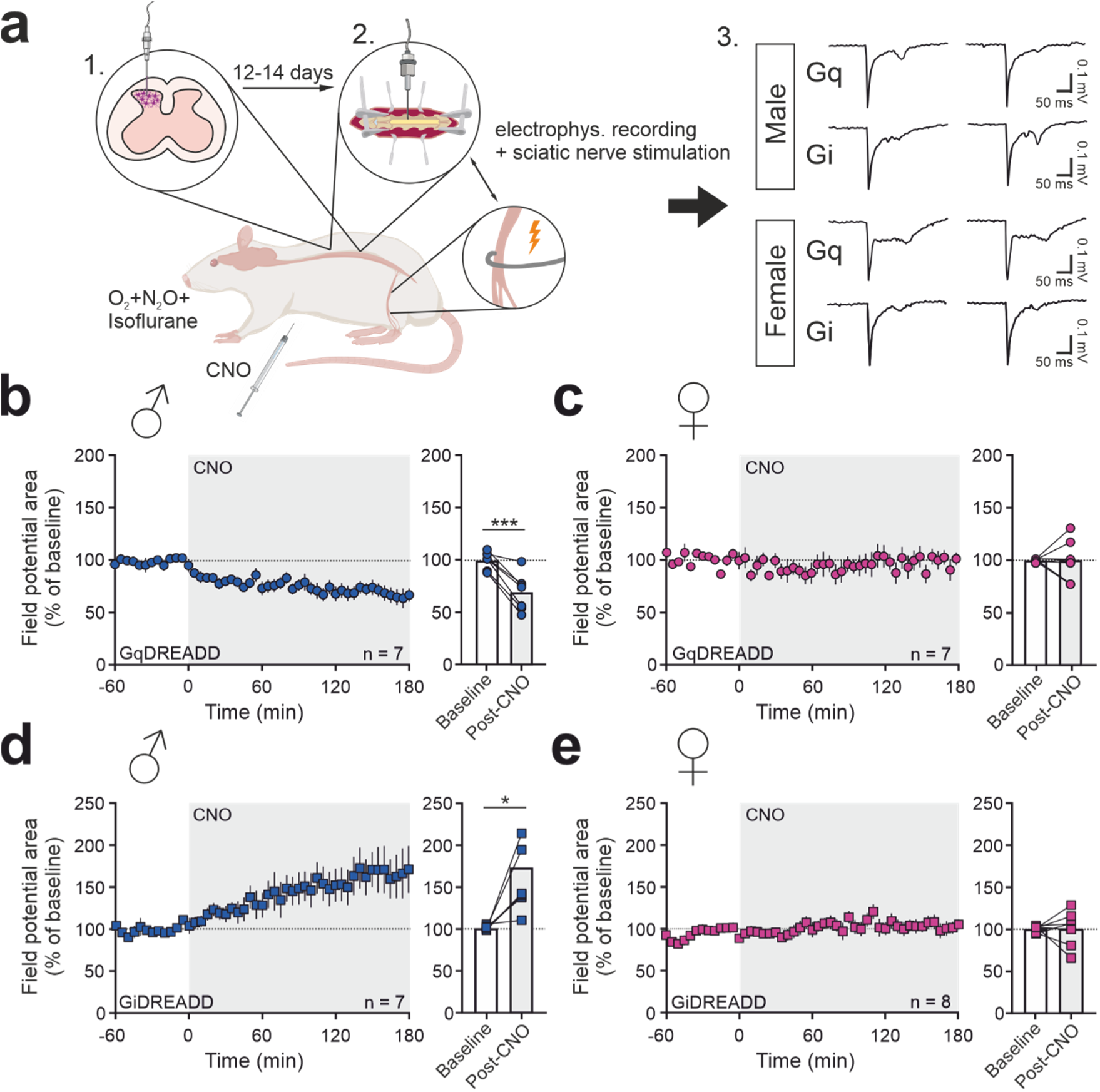
Distinct astrocyte signaling pathways bidirectionally modulate nociceptive transmission in a sex-dependent manner. (**a**) C-fiber-evoked field potentials were recorded in vivo from the SDH following electrical stimulation of the sciatic nerve in DREADD-expressing animals. Insets show representative traces. (**b**–**e**) Field potential areas were normalized to baseline (BL; dashed line) and plotted over time. CNO was administered intravenously (3 mgꞏkg^-1^) at time point 0. Bar graphs summarize BL-normalized mean responses for the indicated time periods. (**b**) Activation of astrocytic Gq-DREADDs induced synaptic depression in male rats. (**c**) Activation of astrocytic Gq-DREADDs had no effect in female rats. (**d**) Activation of astrocytic Gi-DREADDs induced synaptic potentiation in male rats. (**e**) Activation of astrocytic Gi-DREADDs had no effect in female rats. Data are presented as mean ± SEM. Paired t-test, *P < 0.05, ***P < 0.001. Details are given in **Supplementary Table 1**.

To test whether this sex-dependent regulation extended to another astrocytic GPCR pathway, we expressed Gi-coupled DREADDs in spinal astrocytes. In males, Gi-DREADD activation robustly potentiated C-fiber-evoked responses, revealing pathway-dependent, bidirectional astrocytic control of synaptic strength (**Fig. 1d**). This effect was again absent in females (**Fig. 1e**).

Because in vivo recordings preserve intact circuitry but limit cellular resolution, we next performed whole-cell patch-clamp recordings in acute spinal cord slices (**Fig. 2a**). In lamina I neurons receiving monosynaptic C-fiber input, bath application of CNO (10 µM) induced sustained synaptic depression in males expressing astrocytic Gq-DREADDs, but not in sham controls (**Fig. 2b** and **Extended Fig. 2b**), consistent with astroSD observed in vivo and previously reported (*11*). The same manipulation had no detectable effect in females (**Fig. 2c** and **Extended Fig. 2c**).

**Fig. 2.**
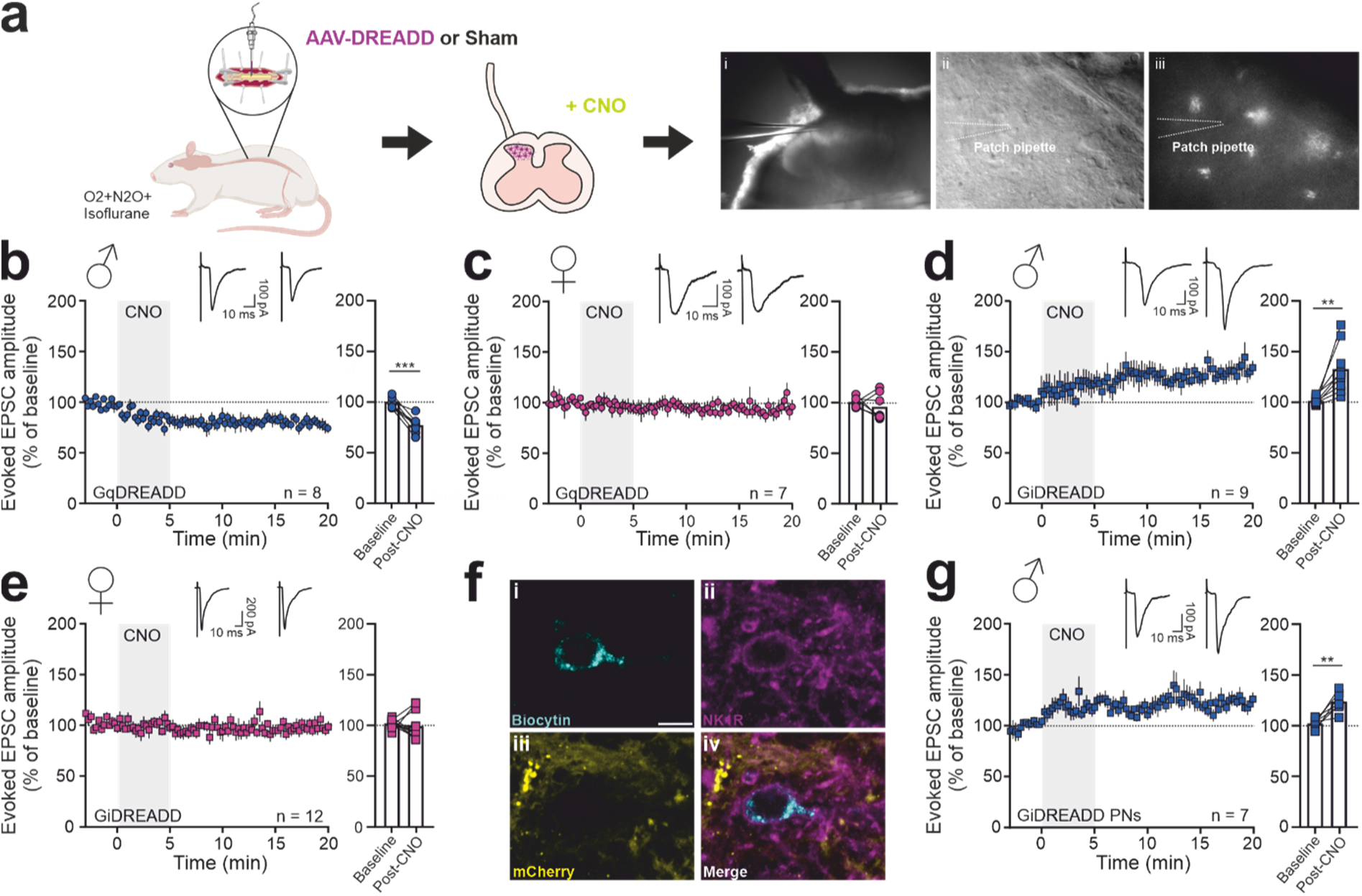
Astrocyte-driven synaptic plasticity is pathway- and sex-dependent at the cellular level. Acute spinal cord slice recordings were performed using dorsal root stimulation and whole-cell patch-clamp recordings from dorsal horn neurons. (**a**) Schematic of the experimental approach. (**b**–**e**, **g**) Amplitudes of C-fiber-evoked EPSCs were normalized to baseline (BL; dashed line) and plotted over time. CNO (10 µM) was bath-applied during the period indicated by the grey shading. Bar graphs summarize BL-normalized mean responses before and after CNO application. (**b**) Activation of astrocytic Gq-DREADDs induced synaptic depression in male rats, but not in females (**c**). (**d**) Activation of astrocytic Gi-DREADDs induced synaptic potentiation in males, but not in females (**e**). (**f**) Representative image of an LPBN-projecting NK1R-positive projection neuron filled with biocytin (i–iv) and surrounded by mCherry-positive DREADD-expressing cells. Individual channels and the merged image are shown. Scale bar, 10 µm. (**g**) Gi-DREADD activation induced synaptic potentiation in LPBN-projecting neurons in males. Data are presented as mean ± SEM. Paired t-test, **P < 0.01; ***P=0.001. Details are given in **Supplementary Table 2**.

In SDH slices from male rats expressing astrocytic Gi-DREADDs, CNO induced rapid, persistent potentiation of evoked EPSCs (**Fig. 2d**), which was absent in females (**Fig. 2e**). Paired-pulse ratio, resting membrane potential, and membrane capacitance were unchanged, arguing against altered release probability or changes in passive membrane properties (**Supplementary Table 2**). To investigate whether astrocyte-induced synaptic plasticity also occurs in neurons conveying nociceptive information to supraspinal targets, we recorded from neurokinin 1 receptor (NK1R)-positive spino-parabrachial projection neurons (**Fig. 2f**). Potentiation in these neurons showed a similar magnitude and time course to that observed in the broader population of lamina I neurons ((*11*); **Fig. 2g**). Therefore, we subsequently performed experiments in unidentified large lamina I neurons with mediolateral orientation.

To explore whether the sex-dependent synaptic effects of astrocytic GPCR activation were correlated with differences in the SD proteome, we performed liquid chromatography-coupled mass spectrometry (LC-MS) in SDH tissue samples. Proteomic datasets showed high technical consistency across biological replicates (**Extended Data Fig. 3a-d**). Importantly, mCherry levels were largely comparable across experimental groups, in particular across sex-matched comparators, suggesting similar viral transduction efficiency (**Extended Data Fig. 3e**). In control conditions (i.e. sham animals), principal component analysis and protein abundance profiles showed overall similarity in male and female control samples, with only two protein groups, Eif2s3 and Nlgn, being significantly altered (**Extended Data Fig. 3f, g**). These data are in agreement with previous reports of modest sexual dimorphism in related sensory tissues such as the dorsal root ganglion (*12*, *13*). In addition, no statistically significant sex differences were detected upon Gq- or Gi-DREADD activation (**Extended Data Fig. 4a, b**). Similarly, we only detected very few proteome changes when comparing DREADD activation with the corresponding controls within each sex (**Extended Data Fig. 4c-f**).

### Astrocyte activation induces mechanical hypersensitivity despite sex-dependent synaptic effects

We next asked whether the pronounced sex differences observed at the level of nociceptive synaptic transmission were reflected in changes in evoked mechanical sensitivity (**Fig. 3a**). Consistent with our previous findings in males (*11*), activation of astrocytic Gq-DREADDs induced transient mechanical hypersensitivity in female animals (**Fig. 3b**). Likewise, activation of astrocytic Gi-DREADDs produced robust mechanical hypersensitivity in both sexes (**Fig. 3c, d**). Thus, females developed mechanical hypersensitivity despite no detectable astrocyte-driven synaptic plasticity, indicating that behavioral hypersensitivity can occur even when astrocytic GPCR activation produces divergent effects on nociceptive synaptic transmission.

**Fig. 3.**
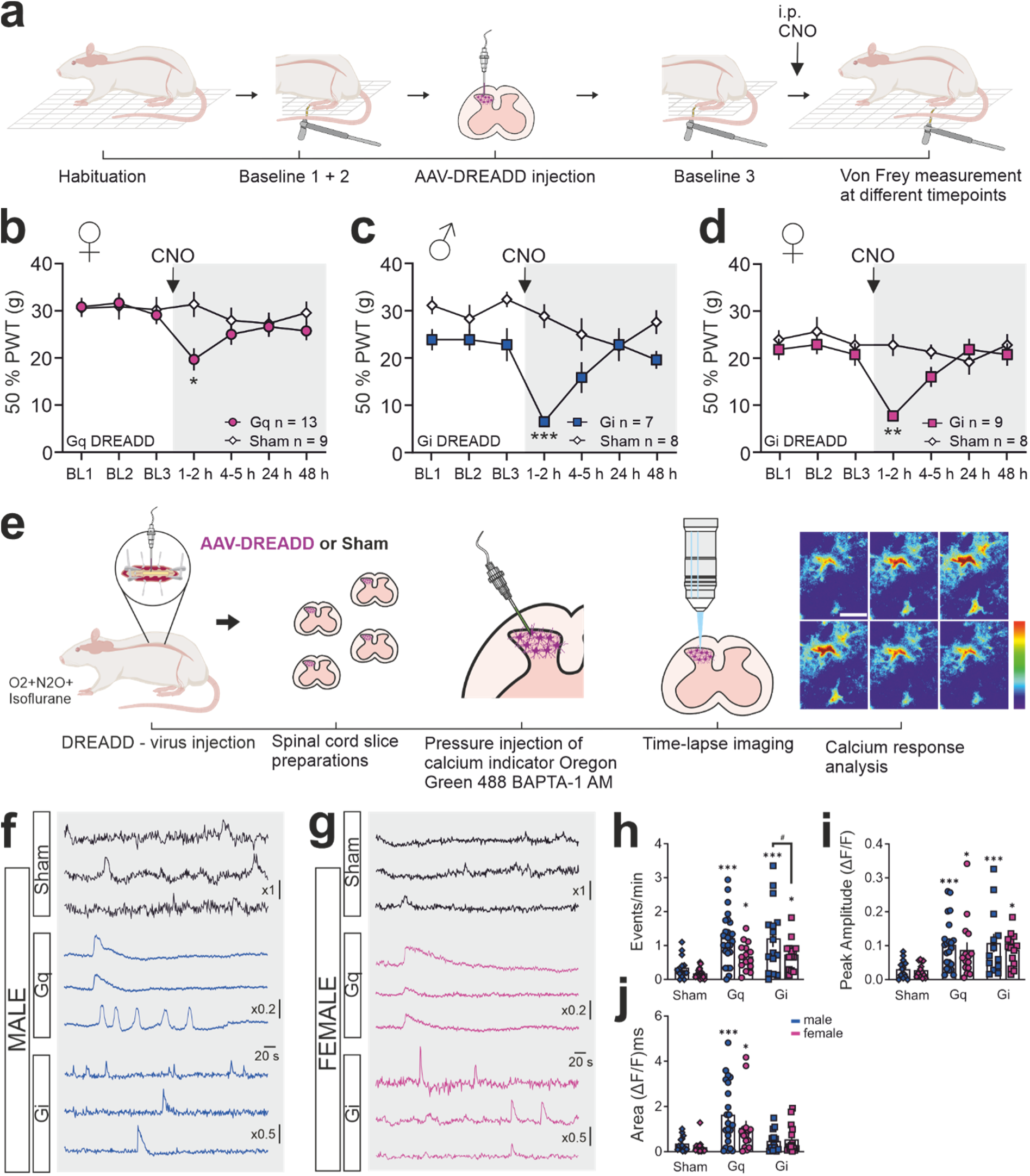
Astrocytic GPCR activation induces mechanical hypersensitivity and engages astrocytic calcium signaling. (**a**) Experimental timeline. Baseline paw withdrawal thresholds (PWTs) were assessed before DREADD injection and again prior to CNO administration. Mechanical sensitivity was measured at the indicated time points following CNO injection. (**b**– **d**) Mechanical withdrawal thresholds in female Gq-DREADD (**b**), male Gi-DREADD (**C**), and female Gi-DREADD (**d**) animals. CNO induced a transient reduction in withdrawal thresholds in all groups, indicating mechanical hypersensitivity. Data are presented as mean ± SEM. Two-way repeated-measures ANOVA, *P < 0.05; **P < 0.01; ***P < 0.001. (**e**) Schematic of the calcium imaging workflow. Scale bar, 10 µm. (**f**, **g**) Representative calcium traces from male and female animals. Beginning of the grey area indicates CNO application. (**h**–**j**) Quantification of calcium event frequency (**h**), peak amplitude (**i**), and event area (**j**). Individual symbols represent individual cells. Data are presented as mean ± SEM. Two-way ANOVA; Asterisks indicate comparisons with the respective same-sex sham controls, and hash symbols indicate comparisons between sexes within the same treatment group. *P < 0.05; ***P < 0.001; #P < 0.05. Additional statistical details are provided in **Supplementary Table 3**.

### Astrocytic GPCR activation engages calcium signaling in both sexes despite divergent synaptic outcomes

To investigate whether the divergent synaptic outcomes could be related to differences in astrocyte activation, we examined calcium dynamics following Gq- and Gi-DREADD stimulation in males and females (**Fig. 3e and Supplementary Table 3**). Representative recordings suggested pathway-dependent differences in the temporal pattern of calcium activity, with Gq-DREADD activation often producing broader, more sustained calcium elevations, whereas Gi-DREADD activation was associated with more discrete calcium events (**Fig. 3f, g**).

No significant sex differences were detected under sham conditions for any of the calcium parameters analyzed (**Fig. 3h-j**). Compared with the corresponding same-sex sham controls, Gq-DREADD activation increased calcium event frequency, peak amplitude, and event area in both males and females. Event frequency was higher in males than in females following Gq-DREADD activation, whereas peak amplitude and event area did not differ between sexes (**Fig. 3h-j**).

Gi-DREADD activation likewise altered astrocytic calcium signaling in both sexes. Compared with the corresponding same-sex sham controls, Gi-DREADD activation increased calcium event frequency and peak amplitude in both males and females, whereas event area was not significantly altered in either sex. No significant sex differences were detected following Gi-DREADD activation for any of the calcium parameters analyzed (**Fig. 3h-j**). Together, these findings demonstrate that both Gq- and Gi-DREADDs functionally engage astrocytic calcium signaling in males and females. While event frequency differed between males and females following Gq-DREADD activation, DREADD-induced changes in calcium signaling were readily detectable in both sexes. Thus, the absence of detectable synaptic plasticity in females occurred despite functional engagement of astrocytic Gq- and Gi-coupled signaling.

To characterize the extracellular cytokine milieu following astrocytic GPCR activation, we performed multiplex cytokine measurements in supernatants collected from whole SDH slices after CNO application. Multiple cytokines were detected following both Gq- and Gi-DREADD activation in males and females (**Extended Data Fig. 5**). Direct comparisons of individual fold changes relative to the corresponding controls revealed no statistically significant differences between males and females within either DREADD condition (**Supplementary Table 8**).

### Hormonal status regulates astrocytogenic synaptic plasticity

We next returned to the striking sex difference observed in astrocyte-induced synaptic plasticity, with Gq- and Gi-DREADD activation inducing synaptic depression and potentiation, respectively, in males but neither form of plasticity in females. To investigate whether gonadal hormone status contributes to this differential responsiveness, we examined the effects of gonadectomy in male and female rats using in vivo electrophysiology (**Fig. 4a**). In gonadectomized males, systemic activation of either Gq- or Gi-DREADDs failed to induce significant synaptic plasticity at the group level (**Fig. 4b, c**), suggesting that gonadal hormone status contributes to astrocyte-driven synaptic modulation in males. Ovariectomy alone did not enable significant astrocyte-driven synaptic plasticity in females (**Fig. 4d, e**).

**Fig. 4.**
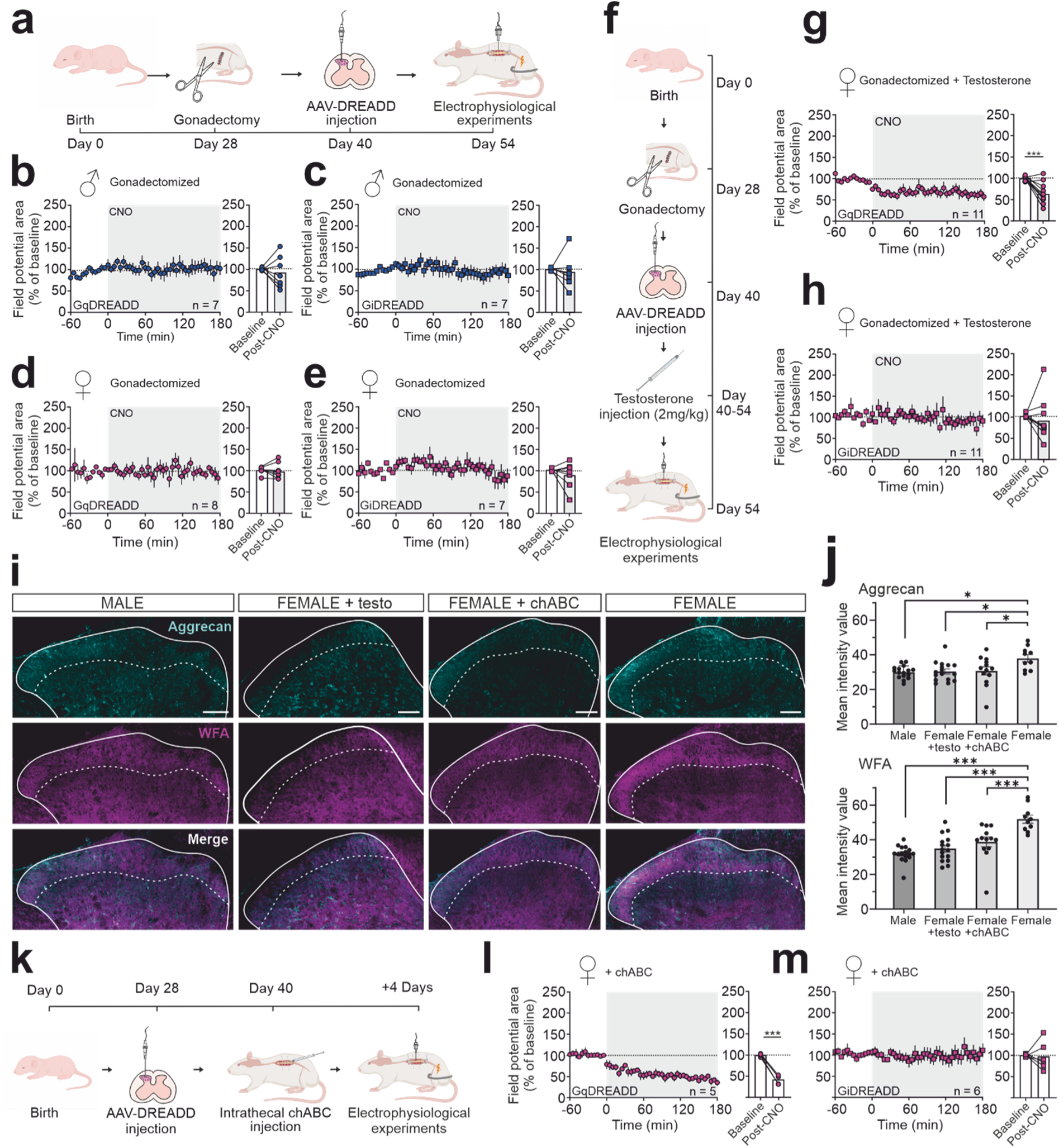
Gonadal hormone status and extracellular matrix organization regulate astrocyte-driven synaptic plasticity. (**a**) Experimental timeline for gonadectomy, AAV-DREADD injection, and in vivo electrophysiological recordings. (**b–e**) Time courses of normalized C-fiber-evoked field potential areas following Gq- (**b**, **d**) or Gi-DREADD (**c**, **e**) activation in gonadectomized male (**b**, **c**) and female (**d**, **e**) rats. (**f**) Experimental timeline for testosterone treatment in ovariectomized females. (**g**, **h**) Normalized C-fiber-evoked field potential areas following Gq- (**g**) or Gi-DREADD (**h**) activation in testosterone-treated ovariectomized females. (**i**) Representative images of aggrecan (cyan) and WFA (magenta) labeling in the spinal dorsal horn of male rats, testosterone-treated females, ChABC-treated females, and untreated females. Scale bar, 100 µm. (**j**) Quantification of aggrecan and WFA fluorescence intensity in spinal dorsal horn sections. (**k**) Experimental timeline for ChABC treatment and in vivo electrophysiological recordings. (**l**, **m**) Normalized C-fiber-evoked field potential areas following Gq- (**l**) or Gi-DREADD (**m**) activation in ChABC-treated females. Field potential areas are expressed as percentages of baseline (BL; dashed line), with CNO administration (3 mgꞏkg⁻¹, i.v.) at time 0 (grey shading). Bar graphs summarize mean responses during the indicated baseline and post-CNO periods. Data are presented as mean ± SEM. Statistical comparisons were performed using one-way ANOVA for fluorescence intensity measurements and paired t-tests for electrophysiological data. *P < 0.05; ***P < 0.001. Additional statistical details are provided in **Supplementary Table 4**.

The loss of astrocyte-driven plasticity after gonadectomy in males prompted us to examine androgenic signaling more directly. Immunostaining revealed androgen receptor immunoreactivity in both astrocytes and neurons in the SDH of male and female rats **(Extended Data Fig. 6a-d**), consistent with multiple potential cellular targets of androgenic signaling within the dorsal horn. We next investigated whether testosterone supplementation could enable astrocyte-driven plasticity in ovariectomized females, allowing us to examine its effects in the absence of endogenous ovarian hormone production. Following two weeks of testosterone treatment, Gq-DREADD activation induced robust astroSD resembling that observed in intact males (**Fig. 4f, g**). In contrast, Gi-DREADD-induced potentiation remained absent in testosterone-treated ovariectomized females (**Fig. 4h**). Notably, testosterone treatment also enabled Gq-dependent astroSD in non-ovariectomized females, indicating that removal of ovarian hormones is not required for this effect (**Extended Data Fig. 6e**).

Serum testosterone concentrations exceeded those measured in intact males, consistent with blood collection shortly after the final injection and confirming systemic androgen exposure (**Extended Data Fig. 6f**).

Together, these findings identify testosterone treatment as a pathway-specific regulator of astrocyte-mediated synaptic plasticity in the SDH. Testosterone restored Gq-dependent astrocytic modulation in females, while Gi-dependent potentiation remained absent in testosterone-treated ovariectomized females, indicating that hormonal state contributes to the conditions under which astrocytic activation can engage nociceptive synaptic plasticity.

### Extracellular matrix organization contributes to sex-dependent astrocyte-driven plasticity

Given the established role of the ECM in regulating synaptic stability and plasticity (*14*, *15*), we next investigated whether sex-dependent differences in ECM organization might contribute to differential susceptibility to astrocyte-driven plasticity and whether testosterone treatment influences these structural features.

Perineuronal nets (PNNs) regulate synaptic remodeling and glial-synaptic interactions (*16*, *17*), are enriched around nociceptive neurons in the SDH (*18*, *19*), and can be influenced by testosterone in other neural circuits (*20*). We therefore assessed PNN organization across sexes using aggrecan and Wisteria floribunda agglutinin (WFA) labeling. PNN labeling was increased in females compared with males throughout the SDH (**Fig. 4i, j**). Notably, testosterone treatment in females reduced aggrecan and WFA labeling to levels comparable to those observed in males and significantly below those of untreated females (**Fig. 4i, j**). Thus, testosterone exposure shifted PNN labeling toward the male phenotype in parallel with restoration of Gq-dependent astrocytogenic plasticity.

To determine whether ECM organization functionally contributes to the differential susceptibility to astrocyte-driven plasticity, we investigated the effects of enzymatic ECM degradation using chondroitinase ABC (ChABC) in females. Histological analysis confirmed a significant reduction in PNN labeling in laminae I and II of the spinal dorsal horn following ChABC treatment compared with untreated females (**Fig. 4i, j**). Using the same treatment protocol, female rats received an intrathecal injection of ChABC four days before electrophysiological recordings (**Fig. 4k**) (*21–23*). Following ChABC treatment, CNO administration induced Gq-dependent synaptic depression, whereas Gi-dependent synaptic potentiation was not restored at the group level in females (**Fig. 4l, m**). These findings indicate that ChABC-sensitive ECM components contribute to the regulation of astrocyte-driven synaptic plasticity and suggest that Gq- and Gi-dependent plasticity are subject to at least partially distinct permissive conditions.

### NMDA receptor signaling differentially regulates astrocyte-driven synaptic plasticity

Having established that hormonal and extracellular factors influence whether astrocyte-driven synaptic plasticity can be expressed, we next examined the neuronal signaling mechanisms underlying Gi-dependent potentiation in males. Chelation of postsynaptic calcium with intracellular BAPTA prevented potentiation (**Fig. 5a**), indicating a requirement for postsynaptic calcium signaling. Intracellular GDPβS likewise abolished potentiation (**Fig. 5b**), demonstrating that postsynaptic G-protein-dependent signaling is necessary for this form of plasticity.

**Fig. 5.**
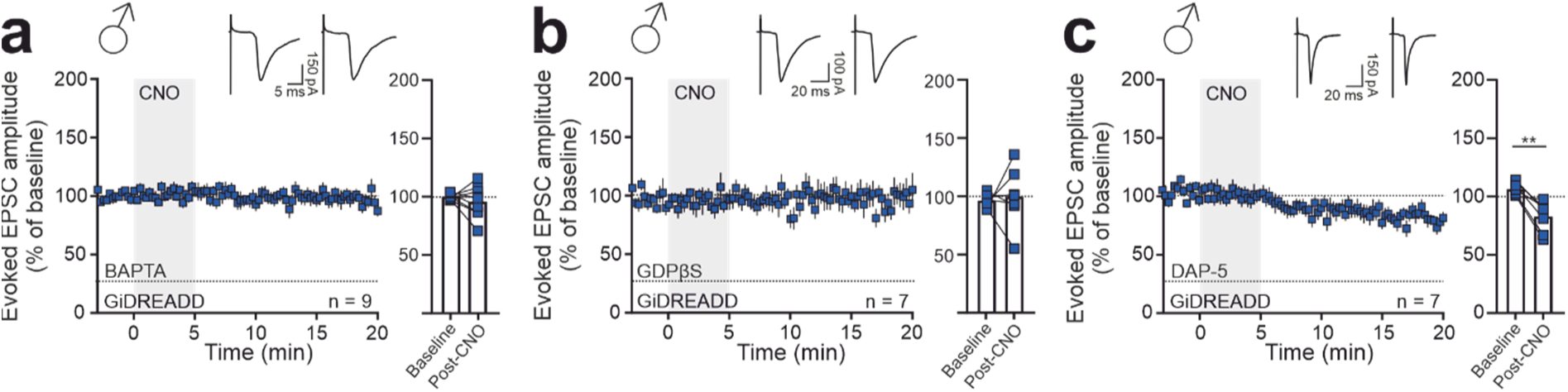
Postsynaptic signaling mechanisms underlying Gi-DREADD-induced synaptic plasticity. Acute spinal cord slice recordings were performed using dorsal root stimulation and whole-cell patch-clamp recordings from dorsal horn neurons. (**a–c**) Time courses of normalized C-fiber-evoked EPSC amplitudes following Gi-DREADD activation in the presence of intracellular BAPTA (**a**), intracellular GDPβS (**b**), or bath-applied D-AP5 (**c**). Representative EPSC traces are shown above each panel. CNO (10 µM) was bath-applied during the period indicated by grey shading. EPSC amplitudes were normalized to baseline (BL; dashed line), and bar graphs summarize mean responses during the indicated baseline and post-CNO periods. Data are presented as mean ± SEM. Paired t-test, **P < 0.01. Additional statistical details are provided in **Supplementary Table 5**.

We then tested the contribution of NMDARs. Bath application of D-AP5 not only blocked potentiation but reversed its polarity, revealing a robust synaptic depression (**Fig. 5c**). Thus, Gi-dependent potentiation requires postsynaptic calcium and G-protein-dependent signaling, as well as NMDAR activation. Notably, NMDAR blockade has distinct consequences for the two forms of astrocyte-driven plasticity: whereas it prevents Gq-induced astroSD, it reveals an opposing depression following Gi-DREADD activation. These findings demonstrate that NMDAR blockade fundamentally alters the synaptic outcome of Gi-DREADD activation, converting the observed potentiation into depression.

## DISCUSSION

Selective activation of distinct astrocytic GPCR pathways revealed a pronounced sex dependence in the ability of spinal astrocytes to engage acute nociceptive synaptic plasticity. In males, Gq- and Gi-DREADD activation induced synaptic depression and potentiation, respectively, whereas females showed no detectable synaptic modification despite preserved astrocytic activation and mechanical hypersensitivity **(Figs. 1, 2 and 3)**. Importantly, hormonal manipulations demonstrated that this difference is not fixed: gonadectomy abolished astrocyte-driven plasticity in males, whereas testosterone treatment was sufficient to restore Gq-dependent synaptic depression in females (**Fig. 4a-h**). In parallel, females exhibited more pronounced perineuronal net labeling than males, which was reduced by testosterone treatment. Consistent with these findings, enzymatic degradation of the ECM using ChABC restored Gq-dependent synaptic depression in females (**Fig. 4i–m**). Neither testosterone treatment nor ECM disruption consistently restored Gi-dependent potentiation in females **(Fig. 4h, m)**. Together, these findings support a model in which sex-dependent susceptibility to astrocyte-driven synaptic plasticity is shaped by hormonal state and ECM organization.

The absence of detectable synaptic plasticity in females cannot readily be attributed to insufficient astrocyte activation. Both Gq- and Gi-coupled DREADDs elicited clear changes in astrocytic calcium signaling in males and females (**Fig. 3f-j**). Gq activation increased calcium event frequency, peak amplitude, and event area in both sexes relative to the corresponding sham controls, whereas Gi activation increased event frequency and peak amplitude. Baseline calcium parameters did not differ between sexes, and the only significant sex difference following DREADD activation was a higher event frequency in males after Gq activation. Thus, female astrocytes were clearly responsive to both GPCR manipulations despite the absence of detectable synaptic plasticity.

Beyond these sex comparisons, representative calcium recordings suggested differences in the temporal characteristics of Gq- and Gi-induced responses, with more sustained calcium elevations following Gq activation and more discrete events following Gi activation **(Fig. 3f, g)**. Such differences could potentially influence the nature or dynamics of extracellular signaling and may be relevant to the opposing forms of synaptic plasticity observed in males (*24*, *25*).

The presence of robust astrocytic calcium responses in females despite the absence of detectable synaptic plasticity suggests that the relationship between astrocytic calcium signaling and synaptic modification depends on the broader cellular and extracellular context. Astrocytic calcium signaling can be associated with the release or regulation of extracellular signaling molecules, which may act on multiple components of the dorsal horn network, including neurons, microglia, and neighboring astrocytes (*26*). Thus, the synaptic consequences of astrocyte activation may depend on how these signals are integrated within the surrounding network, potentially contributing to the sex-dependent susceptibility of nociceptive synapses to astrocyte-driven plasticity.

The sex-dependent regulation of astrocyte-driven synaptic plasticity was not reflected in the behavioral response to astrocyte activation (**Fig. 3a-d)**. Despite the absence of detectable plasticity at the C-fiber synapses examined here, females developed mechanical hypersensitivity following both Gq- and Gi-DREADD activation, comparable to males. This indicates that the forms of synaptic plasticity identified in this study are not universally required for astrocyte-induced mechanical hypersensitivity. In males, Gq- and Gi-DREADD activation induced synaptic depression and potentiation, respectively, yet both manipulations produced mechanical hypersensitivity (*11*). Our previous work further demonstrated that Gq-induced synaptic depression can be converted into potentiation through additional A-fiber stimulation in a microglia-dependent manner, while blockade of afferent activity prevented the development of mechanical hypersensitivity (*11*). Together, these observations highlight that astrocyte activation can produce similar behavioral outcomes despite sex-dependent differences in the expression and direction of synaptic plasticity.

To identify factors contributing to the sex-dependent susceptibility to astrocyte-driven synaptic plasticity, we next examined the contribution of gonadal hormone status. Gonadectomy abolished both forms of astrocyte-driven plasticity in males, whereas testosterone treatment of ovariectomized females selectively restored Gq-dependent synaptic depression but did not consistently restore Gi-dependent potentiation **(Fig. 4a-h)**. Importantly, testosterone treatment also enabled Gq-dependent synaptic depression in non-ovariectomized females, demonstrating that prior removal of ovarian hormones is not required for this effect (**Extended Data Fig. 6e**). This pathway specificity suggests that testosterone does not act as a general permissive factor for astrocyte-driven synaptic plasticity, but that the hormonal conditions permitting synaptic modification differ between the Gq- and Gi-dependent pathways.

Androgen receptor immunoreactivity in astrocyte-marker-positive cells raises the possibility that astrocytes may be directly responsive to androgenic signaling. However, androgen receptor expression in the dorsal horn is not restricted to astrocytes (**Extended Data Fig. 6a-d**), and androgen-dependent modulation of astrocyte-driven plasticity may arise through direct and indirect effects on multiple cellular and extracellular components of the dorsal horn environment (*20*, *27*, *28*).

Among the factors that may influence the susceptibility of spinal nociceptive circuits to astrocyte-driven plasticity, the ECM emerged as a particularly interesting candidate (*15*, *16*, *29*). PNN labeling was markedly higher in females than in males across the SDH (**Fig. 4i, j**), revealing a pronounced sex difference in PNN-associated ECM organization within nociceptive circuits. Testosterone treatment reduced WFA and aggrecan labeling in females toward levels observed in males, in parallel with the restoration of Gq-dependent synaptic depression in gonadectomized females **(Fig. 4g)**. Importantly, ChABC treatment provided functional evidence that the ECM contributes to the susceptibility of nociceptive synapses to astrocyte-driven plasticity. The selective restoration of Gq-dependent depression, but not Gi-dependent potentiation, highlights the pathway-dependent nature of this regulation **(Fig. 4i-m)**. Such differential effects are consistent with previous studies demonstrating that enzymatic ECM degradation can selectively affect distinct forms of synaptic plasticity (*30*, *31*).

PNNs and other ECM components can influence receptor mobility, synaptic stability, extracellular diffusion, and the accessibility of signaling molecules to neuronal membranes (*17*). The more prominent PNN labeling observed in females may therefore reflect extracellular conditions that influence the susceptibility of nociceptive synapses to astrocyte-driven plasticity, even when astrocytes themselves respond to GPCR activation. Although chondroitinase also acts on chondroitin sulfate-containing ECM components beyond classical PNNs (*32*), the functional effects of ChABC, together with the sex-dependent differences in PNN labeling, support a contribution of ECM organization to the sex-dependent regulation of astrocyte-driven plasticity.

Notably, neither testosterone treatment nor enzymatic degradation of the ECM enabled Gi-dependent potentiation in females, whereas both manipulations permitted Gq-dependent synaptic depression. These findings suggest that the mechanisms governing astrocyte-driven plasticity are pathway-specific, with additional or distinct conditions required for Gi-dependent potentiation that are not overcome by either manipulation.

Interestingly, the pronounced baseline sex differences in PNN labeling were not accompanied by statistically significant differences in the abundance of ECM-associated proteins detected by bulk SDH proteomics. However, bulk proteomics assesses overall protein abundance and lacks the spatial and cellular resolution to detect potential changes restricted to specific SDH laminae or cell populations. In contrast, histological labeling reveals the spatial distribution and organization of ECM components around individual neurons. Thus, differences in PNN organization may occur without detectable changes in overall ECM protein abundance, highlighting the importance of spatially resolved analyses for understanding sex-dependent differences in the extracellular environment.

Complementary cytokine measurements identified multiple extracellular cytokines in SDH slice supernatants following Gq-and Gi-DREADD activation in both sexes (**Extended Data Fig. 5**). However, no consistent cytokine profile emerged that could account for the sex-dependent differences in synaptic plasticity. As these measurements were obtained from whole SDH slices, the cellular origin of the detected cytokines cannot be determined.

The two forms of astrocyte-driven plasticity also differed in their neuronal signaling requirements. In males, Gq-DREADD activation induced astroSD, consistent with our previous findings (*11*), whereas Gi-DREADD activation induced potentiation of C-fiber synaptic strength. Gi-dependent potentiation required postsynaptic calcium and neuronal G-protein signaling, and NMDAR blockade prevented potentiation while revealing an opposing depression. This contrasts with Gq-dependent astroSD, in which NMDAR blockade prevented depression without reversing its direction (*11*). Together, these findings highlight the pathway-specific involvement of NMDAR signaling in astrocyte-driven synaptic plasticity, without implying that the underlying mechanisms of depression are shared.

The ability of Gi-coupled astrocytic signaling to induce potentiation is also relevant to the interpretation of astrocytic GPCR manipulation. Gi-coupled signaling is often viewed as functionally inhibitory, largely based on neuronal signaling frameworks (*33–35*). In astrocytes, however, Gi-GPCR activation can evoke intracellular calcium signaling rather than simply suppress cellular activity (*36–42*), consistent with the increased calcium event frequency and peak amplitude observed following Gi-DREADD activation in both sexes here. Thus, Gq- and Gi-coupled GPCR activation should be considered as engagement of distinct astrocytic signaling pathways rather than straightforward activation and inhibition of astrocyte function. Although chemogenetic approaches allow these pathways to be investigated selectively, they do not fully reproduce the spatial and temporal dynamics of endogenous astrocytic signaling. Future studies will be needed to establish their contributions to nociceptive synaptic plasticity under physiological and pathological conditions.

Together, our results support a model in which astrocyte-driven nociceptive synaptic plasticity is gated at multiple levels. Astrocytic GPCR activation elicits calcium responses in both sexes, neuronal signaling mechanisms contribute to distinct forms of synaptic plasticity, and hormonal state and extracellular matrix organization influence whether these modifications can be engaged. Thus, the absence of astrocyte-driven plasticity in females under baseline conditions cannot simply be attributed to insufficient astrocytic activation, but is consistent with differences in the conditions that permit astrocytic signals to modify nociceptive synapses. These conditions may also involve interactions with other cellular populations within the dorsal horn, including microglia, consistent with our previous findings (*11*). Importantly, our findings establish ECM organization as a functional component of the regulation of Gq-dependent plasticity and demonstrate that the susceptibility of nociceptive synapses to astrocyte-driven plasticity can be modified by hormonal and extracellular conditions.

These findings may have broader implications for understanding sex differences in pain mechanisms and developing targeted therapeutic approaches. Comparable mechanical hypersensitivity despite distinct synaptic responses to astrocytic activation suggests that similar nociceptive phenotypes may involve different underlying circuit mechanisms. Furthermore, the dependence of astrocyte-driven plasticity on hormonal and extracellular conditions raises the possibility that interventions targeting glial signaling may have context-dependent effects. Although the clinical implications remain to be determined, our findings suggest that sex-dependent circuit properties and the local neuronal environment should be considered when exploring glia-targeted strategies for pain treatment.

## MATERIALS AND METHODS

### Animals

All experiments were performed on male and female Sprague–Dawley rats. Animals were obtained either from Janvier Labs or from the Medical University of Vienna’s breeding facilities. Rats were group-housed in standard cages under environmentally controlled conditions, including regulated temperature and humidity and a defined light–dark cycle. Food and water were available ad libitum, and animals received appropriate environmental enrichment, including shelter tubes, chew wood, and nesting material. Estrous cycle stage was not monitored in intact females. Female animals were therefore analyzed under mixed estrous-cycle conditions, reflecting sex-dependent effects in the presence of physiological hormonal variability, but not allowing assessment of cycle stage-dependent modulation.

For experiments involving viral vector or tracer injections, rats were injected at 25 – 35 days of age and used for subsequent experiments only after an expression period of at least 15 days. All other experimental procedures were performed on adult animals (7 – 8 weeks old). A subset of animals underwent gonadectomy before injections, which was performed at Janvier Labs.

All experimental procedures were carried out by trained personnel and conducted in accordance with national and European legislation governing the use of animals for scientific purposes, as well as the regulations of the Austrian Ministry for Science. Experiments were approved by the relevant institutional and governmental authorities and complied with all requirements for animal welfare, refinement, and minimization of animal burden.

### Intraspinal injections

For viral delivery, animals were anaesthetized with isoflurane (5 vol% for induction, 3.5 vol% for maintenance) and ventilated via a face mask. Respiratory rate was maintained between 80 and 90 breaths per minute. Body temperature was continuously monitored and kept between 35 and 37 °C using a feedback-controlled heating pad. Ophthalmic ointment was applied to prevent corneal drying, and buprenorphine (0.05 mgꞏkg^-^¹, s.c.) was administered for analgesia.

After loss of the tail pinch reflex, a midline skin incision was made, and the dorsal musculature was bluntly dissected. A unilateral hemilaminectomy was performed to expose the L4/L5 segment of the lumbar spinal cord. The vertebral column was stabilized using a clamp attached to a custom–built frame. Viral injections were performed using borosilicate glass pipettes with a tip diameter of 30 to 40 µm. The pipette was positioned approximately 100 µm lateral to the central vein and advanced to a depth of about 200 µm below the tissue surface.

A total of 500 nl of ssAAV-9/2-hGFAP-hM3D(Gq)_mCherry-WPRE-hGHp(A), ssAAV-9/2-hGFAP-hM4D(Gi)_mCherry-WPRE-hGHp(A), or ssAAV-9/2-hGFAP-mCherry-WPRE-hGHp(A) (as control; all obtained from the Viral Vector Facility, Neuroscience Center Zurich, Switzerland; titer 6 x 10¹² vg/ml) was injected using a motorized microinjection pump (Nanoject III, Drummond Scientific, USA) at 50 nlꞏmin^-^¹. A second injection was placed 1 to 2 mm caudal to the first site. The pipette was left in place for an additional 10 minutes after each injection to allow diffusion of the viral solution.

After completion of the injections, animals were released from the vertebral clamp, isoflurane concentration was reduced to 2 vol%, and the wound was sutured and treated with antibiotic ointment. Rats received subcutaneous injections of carprofen (4 mgꞏkg^-^¹) for postoperative analgesia, before they were housed individually and monitored until they fully recovered from anesthesia. Animals were reunited with their littermates one day after surgery and subsequently housed in groups of three.

### Supraspinal injections

For supraspinal injections, projection neurons were retrogradely labelled as described previously (*11*, *43*). Rats were deeply anaesthetized with isoflurane (3.5 vol% for maintenance) in combination with ketamine and xylazine (50 and 4 mgꞏkg^-^¹, i.p., respectively). Animals were positioned in a stereotaxic frame, and a burr hole was drilled 1.8 mm lateral and 0.5 mm caudal to the apex of the lambdoid suture. A micro syringe was inserted at a 7° angle, and cholera toxin subunit B conjugated to Alexa Fluor 555 (CTB-555, Invitrogen C34776; 200 nl of a 1% solution) was infused into the parabrachial nucleus at a rate of 0.1 µlꞏmin^-^¹. After the injection, the wound was disinfected, and the skin was sutured. Carprofen (4 mgꞏkg^-^¹, s.c.) was administered for postoperative analgesia. Only animals with correctly targeted injections, verified according to the Paxinos and Watson atlas (*44*), were included in the analysis. Robust labelling of spinal neurons was typically observed three to five days after tracer injection.

### Preparation of spinal cord slices and electrophysiological recordings in vitro

Fifteen days after the spinal injections, or up to five days following supraspinal injections, animals were killed under deep isoflurane anesthesia and their spinal cord removed to prepare 500 µm thick transverse spinal cord slices with the dorsal roots attached using a vibratome. Slices were first incubated for 30 minutes at 37 °C and subsequently maintained at room temperature (approximately 21 °C) in continuously oxygenated incubation solution containing (in mM): 95 NaCl, 1.8 KCl, 1.2 KH₂PO₄, 0.5 CaCl₂, 7 MgSO₄, 26 NaHCO₃, 15 glucose, and 50 sucrose (pH 7.4; osmolarity 305 – 320 mOsmolꞏl^-^¹). For recordings, slices were transferred to a recording chamber and superfused at 3 – 4 mlꞏmin^-^¹ with oxygenated recording solution. The recording solution differed from the incubation solution in that it contained (in mM): 127 NaCl, 2.4 CaCl_2_, 1.3 MgSO_4_, and no sucrose. All recordings were performed at room temperature.

Successful viral transduction was confirmed by visual detection of mCherry fluorescence. Neurons located in lamina I within regions of mCherry expression were visualized using Dodt infrared contrast optics with a 40x water-immersion objective (NA 0.80) on an upright Olympus BX50WI microscope.

Whole-cell patch-clamp recordings were obtained using glass micropipettes (2 – 4 MΩ). For current-clamp recordings and measurements of evoked excitatory postsynaptic currents (eEPSCs), pipettes were filled with an intracellular solution containing (in mM): 120 potassium gluconate, 20 KCl, 2 MgCl₂, 20 HEPES (4-(2-hydroxyethyl)-1-piperazineethanesulfonic acid, Sigma Aldrich, H-3375), 0.5 Na-GTP (guanosine 5′-triphosphate sodium salt hydrate, Sigma Aldrich, G8877), 0.5 Na_4_-EGTA (Na_4_-ethylene glycol-bis(β-aminoethyl ether)-N,N,N′,N′-tetraacetic acid, Sigma Aldrich, E-8145), 2 Na₂-ATP (Na₂-adenosine triphosphate, Sigma Aldrich, A-2383), and 7.5 Na₂-phosphocreatine x2 H_2_O (pH 7.4, adjusted with KOH; 310 mOsmꞏkg^-^¹, Sigma Aldrich, P-7936). Biocytin (2 mgꞏml^-^¹, Biotium, 90055) was included for post hoc morphological identification of recorded neurons. Resting membrane potential was measured immediately after establishing whole-cell access. Membrane capacitance was determined by averaging responses to twenty 100-ms hyperpolarizing voltage steps from −70 to −80 mV and integrating the capacitive transient.

eEPSCs were recorded at a holding potential of −70 mV using an Axopatch 700B amplifier and pClamp 10 software. Signals were low-pass filtered at 2-10 kHz, digitized at 20 kHz, and analyzed offline. Liquid junction potentials were not corrected. Following each recording, images of the neuron were acquired, and the distance from the soma to the ventral border of the white matter was measured using ImageJ.

eEPSCs were elicited by electrical stimulation of dorsal roots via a suction electrode. Stimulation intensity was adjusted to activate C-fiber afferents. Conduction velocity was calculated from the latency between the stimulation artifact and eEPSC onset; fibers with conduction velocities <0.5 m s⁻¹ were classified as C-fibers. Test stimulus intensity was set to twice the threshold required to evoke a C-fiber eEPSC. Monosynaptic responses were identified by constant latency and the absence of failures during repetitive dorsal root stimulation at 1 Hz.

Test stimuli were delivered every 15 s. Paired-pulse stimulation (0.1-ms pulse duration, 300-ms inter-pulse interval) was used to determine paired-pulse ratios. Series resistance was continuously monitored throughout the experiment, and recordings were excluded if series resistance exceeded 25 MΩ or changed by more than 30%.

To activate astrocytic DREADDs, clozapine-N-oxide (CNO, 10 µM, APExBIO, A3317) was added to the recording solution.

Synaptic strength was quantified as the peak amplitude of eEPSCs using Clampfit 10 software (version 10.7.0.3). Responses were normalized to the mean peak amplitude of the final eight sweeps recorded during the 2-minutes period preceding CNO application, which was defined as baseline. Statistical comparisons between time points were performed using paired t-tests in GraphPad Prism 9 (version 9.5.1). Further details of the statistical analysis are provided in **Supplementary Tables 2, 5 and 7**.

### Calcium imaging

Calcium imaging experiments were performed as previously described (*11*). Acute transverse spinal cord slices (500 µm) were prepared from animals injected with Gq-DREADD, Gi-DREADD, or control virus (see above). Slice preparation followed the procedures described above. After sectioning, slices were allowed to recover for at least 30 minutes at 32 °C in oxygenated incubation solution and were subsequently maintained in the same solution at room temperature until use.

For imaging, slices were transferred to a recording chamber and continuously superfused with oxygenated recording solution. Following confirmation of mCherry expression, the calcium indicator Oregon Green^TM^ 488 BAPTA-1 AM (1,2-Bis(2-aminophenoxy)ethane-N,N,N′,N′-tetraacetic acid tetrakis(acetoxymethyl ester), Invitrogen, O6807) was locally delivered into the dorsal horn at a minimum of three distinct sites. Dye loading was performed by pressure injection using glass micropipettes (3 – 4 MΩ) containing HEPES-buffered artificial cerebrospinal fluid supplemented with 353.3 µM Oregon Green^TM^-BAPTA-1 AM, 1.6% Pluronic F-127, and 3.4% dimethyl sulfoxide (DMSO, Sigma Aldrich, D-4540). Slices were then incubated for 30 – 40 minutes to allow for intracellular de-esterification of the dye.

Time-lapse imaging was carried out at room temperature using a confocal microscope (DM6000 CFS, Leica, Germany) equipped with a 20x objective (HCX APO, NA 1.0). Fluorescence images were acquired at a rate of 1 Hz from a single focal plane.

Image processing and analysis were performed offline using Fiji (ImageJ, version 1.54t). Recorded time series were imported using BIO-Formats plugins (release 6.11.0). Lateral drift in the xy plane was corrected using the Multi Drift Correction plugin (version 1.1), while recordings showing detectable z-axis drift were excluded from analysis. Cells positive for Oregon Green BAPTA-1 AM and co-expressing mCherry were selected for analysis as DREADD-expressing astrocytes, based on the independently validated astrocytic specificity of the viral construct (**Extended Data Fig. 1**). Cells positive for Oregon Green^TM^-BAPTA-1 AM and co-expressing mCherry were identified as astrocytes, and Regions of interest were manually defined for these cells. CNO (10 µM) was bath-applied.

Fluorescence traces were extracted and further analyzed using a custom script (https://github.com/ronihogri/Calcium-signal-preprocessing-and-analysis-with-Spike2.git) written in Spike2 (Cambridge Electronic Design, UK; version 9.01 x64). Signal changes were expressed as ΔF/F₀, calculated as (F(t) − F₀) / F₀. To compensate for slow baseline fluctuations, F₀ was determined dynamically using a sliding window of the preceding 30 subthreshold data points, with the detection threshold set to the mean plus two standard deviations of the window. The initial 10 seconds of each recording served as the baseline reference window. Subsequently, traces were high-pass filtered at 0.001 Hz using a Butterworth filter.

Calcium transients were identified as periods in which ΔF/F₀ exceeded the baseline mean by two standard deviations for a minimum duration of 3 seconds. For each detected event, the peak amplitude was quantified. Only responsive cells were included in the analyses.

Statistical comparisons were performed using one-way ANOVA followed by Tukey’s post hoc test. Further details of the statistical analysis are provided in **Supplementary Table 3**.

### Surgery for in vivo recordings

Detailed procedures have been described elsewhere (*11*). Briefly, male and female rats were deeply anaesthetized with a mixture of 5% isoflurane in oxygen/nitrous oxide. Animals were then intubated using a 16-gauge cannula and mechanically ventilated. Anesthesia was maintained with 2.5 – 3% isoflurane throughout the experiment. Body temperature was kept constant using a feedback-controlled heating blanket.

The carotid artery and jugular vein were cannulated, and the sciatic nerve was carefully dissected free, with particular care taken to avoid direct contact with the nerve fibers. A laminectomy was subsequently performed to expose lumbar spinal segments L4 to L6, followed by careful removal of the dura mater.

The spinal column was stabilized using two custom-built clamps. A recording pool was created by suturing the surrounding skin and filling the cavity with agarose to cover all exposed tissue except the spinal cord, which was continuously superfused with artificial cerebrospinal fluid consisting of the following composition (in mM): 135 NaCl, 1.7 KCl, 1.8 CaCl_2_, 10 HEPES, 1 MgCl_2_, pH 7.4 adjusted with KOH, with an osmolarity of 290 mOsmolꞏl^-1^, using a roller pump within a closed circuit system. The sciatic nerve was placed on a silver hook electrode to enable electrical stimulation.

Animals received a continuous intravenous infusion of physiological saline containing pancuronium bromide at a rate of 2 mlꞏh^-1^. Physiological parameters were monitored regularly using arterial blood gas analysis and pulse oximetry to control blood oxygenation, heart rate, and respiratory rate.

### Testosterone treatment

**I**n some experiments, gonadectomized male and female animals were used. Castration or ovariectomy was performed at Janvier Labs, and animals were delivered approximately one week after surgery. At the time of delivery, the animals were approximately 5 weeks old. They were allowed to acclimate in the animal facility for at least 14 days before any further surgical procedures or viral injections were performed.

Where indicated, animals were additionally treated with testosterone. For this purpose, testosterone was administered by daily subcutaneous injection. A solution at 5 mgꞏml^-1^ was used, corresponding to an injection volume of 0.4 mlꞏkg^-1^ body weight. Injections were given once daily between 7 and 9 a.m. for two consecutive weeks to establish a testosterone-rich hormonal milieu.

To assess testosterone levels, blood samples were collected immediately before the start of the electrophysiological recordings via the intra-arterial catheter into coated Eppendorf tubes. After allowing the samples to clot for 30 min at room temperature, they were centrifuged at −4 °C for 10 min at 1200 rpm, and the resulting serum was collected and stored at −80 °C for subsequent analysis.

### Chondroitinase ABC Treatment

In a subset of experiments, chondroitinase ABC (ChABC) was used to enzymatically degrade chondroitin sulfate proteoglycans, including those associated with perineuronal nets (*21–23*). Ten to twelve days after Gq-or Gi-DREADD injection, female rats received an intrathecal injection of ChABC (20 µl, 10 Uꞏml^-^¹) in artificial cerebrospinal fluid (aCSF) supplemented with 60 mM sodium acetate. The enzyme solution was freshly prepared before each experiment. Four days after ChABC administration, animals were either prepared for in vivo electrophysiological recordings or perfused for immunohistochemical analysis.

### In vivo electrophysiology

Glass recording pipettes (3-4 MΩ) were filled with a solution containing (in mM): 135 NaCl, 1.7 KCl, 1.8 CaCl₂, 10 HEPES, 1 MgCl₂, and 0.2% rhodamine B (Sigma Aldrich, R-6626). Field potentials were evoked by electrical stimulation of the sciatic nerve at C-fiber strength and recorded from the superficial laminae of the spinal dorsal horn. A supramaximal test stimulus (0.5 ms, 5 V) was delivered every 5 minutes throughout the recording period. CNO (3 mgꞏkg^-^¹) was administered intravenously (i.v.).

At the end of each experiment, brief pressure was applied to the recording pipette to deposit rhodamine and thereby mark the recording site. Animals were then euthanized by decapitation, and the spinal cord was rapidly removed and flash-frozen in isopentane. Tissue sections were subsequently prepared to verify the location of the recording site. Only animals in which the rhodamine-labelled recording site was located within the superficial dorsal horn and within the region of viral expression were included in the final analysis.

Synaptic strength was quantified as the area of the C-fiber-evoked field potential using Clampfit 10 software. Responses were normalized to the mean area of the final six responses recorded during the 60-min baseline period preceding CNO administration. Statistical comparisons between time points were performed using paired t-tests in GraphPad Prism 9. Further details of the statistical analysis are provided in **Supplementary Tables 1, 4 and 7**.

### Proteomics

#### Tissue preparation

At 10 – 12 days after AAV injection, male and female rats received an i.p. injection of CNO (3 mgꞏkg⁻¹). SDH tissue was collected under deep isoflurane anesthesia 2 h after CNO administration. This sampling time point was selected to coincide with the period of established astrocyte-induced synaptic plasticity in our in vivo electrophysiological experiments and the observed behavioral responses.

The spinal cords were then sectioned into 500 µm transverse slices using a vibratome. Slices were incubated for 30 minutes in oxygenated incubation solution (recipe listed above) at 37 °C and subsequently maintained at room temperature.

Slices were then visually inspected for viral expression, and only those exhibiting sufficient expression in the dorsal horn were included for further experiments. For proteomic measurements, the ipsilateral dorsal horns (based on the side of viral injection) were microdissected from a minimum of five slices with detectable mCherry expression, transferred into 1.5 ml Eppendorf tubes, shock-frozen in liquid nitrogen, and subsequently stored at −80 °C until further analysis.

#### Protein extraction

A volume of 250 µl lysis buffer (2% SDS, 100 mM Tris, 5% glycerol, 10 mM DTT, and 1× protease inhibitor cocktail − cOmplet Mini, Sigma-Aldrich, 4693159001) was added to LoBind tubes containing tissue slices. Samples were vortexed and subsequently sonicated for 10 min using a Bioruptor Pico (Diagenode, Seraing, Belgium) at 4 °C (10 cycles of 30 s ON and 30 s OFF, low frequency). Following sonication, samples were incubated for 10 minutes at 70 °C with shaking at 150 rpm in a thermomixer. To remove cellular debris, samples were centrifuged for 5 minutes at 10000 × g at 70 °C. The supernatant was transferred to a new LoBind tube and stored at −20 °C. To reduce lipid content, acetone precipitation was performed. Five volumes of pre-cooled (−20 °C) 100% acetone were added to each sample, followed by brief inversion to ensure thorough mixing. Where necessary, samples were split into two 2 ml tubes to allow for the 5× volume addition. Protein precipitation was carried out at −20 °C for 3.5 h. Proteins were then pelleted by centrifugation at 14000 × g for 30 min at room temperature. The resulting pellets were washed with 1.5 ml pre-chilled (4 °C) 80% ethanol and centrifuged again under the same conditions. After removal of the ethanol, pellets were air-dried for 20 minutes. Proteins were reconstituted in lysis buffer by incubation at 60 °C for 10 min with shaking at 1500 rpm in a thermomixer and subsequently kept on ice. After a brief centrifugation (1 min at 14000 × g, room temperature), total protein concentration of pooled samples was determined at 280 nm using a NanoPhotometer N60 (Implen, Germany). Samples were stored at −20 °C.

#### Protein digestion

For protein clean-up and digestion, a modified version of the single-pot, solid-phase-enhanced sample preparation (SP3) method from Hughes et al. was used (*45*). Briefly, 10 μl of pre-mixed Sera-Mag SpeedBead beads (Cytiva, Marlborough, MA) were added into a 50 μg protein sample. To initiate binding of proteins to the beads, one volume of absolute ethanol was added immediately, followed by incubation on a Thermomixer (Eppendorf) at 24 °C for 5 minutes with 1000 rpm agitation. The supernatant was removed after 2 minutes resting on a magnetic rack, and the beads were rinsed three times with 500 μl of 80% ethanol. Rinsed beads were reconstituted in a 50 μl digestion buffer (50 mM ammonium bicarbonate, pH 8). Protein digestion was performed with 2 μg of sequencing grade trypsin/Lys-C for 18 hours at 37 °C with 950 rpm agitation. After digestion, acetonitrile was added to each sample to a final concentration of 95%. Mixtures were incubated for 8 min at room temperature and then placed on a magnetic rack for 2 min. The supernatant was discarded, and the beads were rinsed with 900 μL of 100% acetonitrile. The rinsed beads were reconstituted with 30 μl LC-MS grade water to elute the peptides. Peptide concentration was measured in duplicate using NanoPhotometer N60 at 205 nm and samples were stored at −20 °C until LC-MS/MS analysis.

#### Liquid chromatography coupled to mass spectrometry

Nanoflow reversed-phase liquid chromatography (Nano-RPLC) was performed using a NanoElute2 system (Bruker Daltonik, Bremen, Germany) and coupled to a timsTOF HT mass spectrometer (Bruker Daltonik, Bremen, Germany) via a CaptiveSpray ion source (Bruker Daltonik, Bremen, Germany). A total of 450 ng of peptides were loaded for each analysis. The mobile phase consisted of solvent A (100% water, 0.1% formic acid) and solvent B (100% acetonitrile, 0.1% formic acid). Peptides were first loaded onto a C18 trap column (1 mm × 5 mm, Thermo Fisher) before being separated on Aurora™ ULTIMATE analytical column (25 cm × 75 µm) packed with 1.6 µm C18 particles (IonOpticks, Fitzroy, Australia) over a 60-minute gradient. The flow rate was maintained at 250 nl/min, except for the final 7 minutes, where it was increased to 400 nl/min. The gradient of mobile phase B was adjusted as follows: an initial linear increase from 4% to 20% over 35 minutes, followed by a rise to 35% over 17 minutes, and a sharp increase to 85% within 0.5 minutes. The flow rate was then adjusted to 400 nl/min in 0.5 minutes and held for 7 minutes to ensure complete elution of hydrophobic peptides. Data acquisition was performed in data-independent acquisition (DIA) mode using parallel accumulation serial fragmentation (PASEF). Precursors with an m/z range of 350 – 1250 were analyzed across 14 MS/MS ramps in a cycle, with either two or three quadrupole switches per ramp. Each cycle contained 36 MS/MS windows within a 0.7 – 1.40 (1/k₀) ion mobility range, using a fixed isolation window of 25 Da per step. The acquisition time per DIA-PASEF ramp was set to 100 ms, resulting in a total cycle time of approximately 1.59 seconds. Collision energy was ramped linearly from 65 eV at 1/k₀ = 1.6 to 20 eV at 1/k₀ = 0.6.

#### Database searches, quantification, and normalization

DIA-NN (*46*) (version 2.2.0) was used to process DIA-PASEF data in library-free mode on the Austrian Scientific Cluster. A deep learning-based method was used to predict theoretical peptide spectra along with their retention time and ion mobility from a reviewed proteome database of *Rattus norvegicus* from UniProt containing 8212 protein entries (downloaded on 2025-06-26), supplemented with the sequence of the PAmCherry reporter. Trypsin/P was used for in silico digestion with an allowance of a maximum of 2 missed cleavages. Variable modifications on peptides were set to N-terminal methionine excision, methionine oxidation and N-terminal acetylation, while carbamidomethylation of cysteine was a fixed modification. The maximum number of variable modifications on a peptide was set to 2. Peptide length for the search ranged from 7 to 30 amino acids. The m/z ranges were set in accordance with the DIA-PASEF acquisition method to 350–1250 for precursor ions and 100–1700 for fragment ions. Mass accuracy for both MS1 and MS2 was set to automatic determination. Scoring and proteotypicity were set to “Peptidoforms” and “Genes”, respectively, and the option “Protein inference” was unchecked. Match-between-runs (MBR) was enabled for cross-run analysis. RT-dependent cross-run normalization and QuantUMS (high precision) options were selected for quantification (*47*). The DIA-NN search outputs were further processed with the diann R package (version 1.0.1) to calculate the MaxLFQ quantitative intensities for all identified gene groups with q-value ≤ 0.01 as criteria at precursor and gene group levels (*48*). Gene group outputs are also referred to as “proteins” throughout. Repeated injections of a pooled quality-control (QC) sample (n = 4) were quantified in an earlier DIA-NN search of all samples with the same settings but without the PAmCherry sequence; only the QC injections from this search were used, and only for technical quality control.

#### Data pre-processing and quality control

All analyses were performed in R (version 4.4.1) (*49*). MaxLFQ intensities were log_2_-transformed and zero values were treated as missing; missing values were not imputed. Samples were annotated with sex, genotype (control, Gq, Gi) and time point from the sample metadata. Technical reproducibility was assessed from the QC pool injections by the Pearson correlation of log_2_ intensities between injections and by the coefficient of variation (CV) of each gene group quantified in at least 3 of the 4 injections. Within-group biological variability was assessed as the CV of each gene group quantified in at least 3 animals of a sex × genotype group. Expression of the viral construct was confirmed by the intensity of the PAmCherry reporter in each animal. Because spinal cord samples can contain variable amounts of blood, a blood-contamination score was calculated for each sample. Twenty-eight genes encoding plasma proteins (haemoglobins, albumin, fibrinogens, apolipoproteins and other liver-derived plasma proteins) were selected, of which 20 were quantified; plasma proteins with known substantial expression in glial cells (e.g. Agt, C3, C4, Cp, A2m) were deliberately excluded. Log_2_ intensities of each quantified blood protein were standardised across all samples (*z*-scores), and the blood score of a sample was defined as the mean *z*-score of these proteins. Samples with a blood score above 0.8 were excluded (n = 4). The final dataset comprised 26 animals: control males n = 5, control females n = 5, Gq males n = 4, Gq females n = 4, Gi males n = 5 and Gi females n = 3. Sample quality was further assessed by the number of quantified gene groups per sample, sample-to-sample correlation and principal component analysis (PCA).

#### Differential expression analysis and principal component analysis

Differential expression analysis was conducted using the limma package (version 3.62.2) in R (*50*). Proteins quantified in at least 75% of the animals of at least one sex × genotype group were included in the model. A design matrix with one coefficient per sex × genotype group (six groups) was constructed, and a linear model was fitted to the log_2_ intensities of each protein using the lmFit function with all analysed animals. Comparisons were then extracted as linear contrasts (makeContrasts, contrasts.fit); sexes were never pooled. Seven comparisons were tested: females vs males within each genotype (control, Gq and Gi), and Gq vs. control and Gi vs. control within males and within females. Empirical Bayes moderation was applied with the eBayes function (default settings), and statistical significance was assessed using moderated t-tests. To ensure robust quantification, a protein was tested in a given comparison only if it was quantified (non-zero value) in at least 75% of the animals of each of the two groups compared (e.g. 4 of 5, 3 of 4 or 3 of 3 animals); 5268–5426 proteins were tested per comparison (5612 in at least one comparison). P-values were adjusted for multiple testing within each comparison (over the proteins tested in that comparison) using the Benjamini–Hochberg method to control the false discovery rate (FDR), and proteins with an adjusted *P* < 0.05 were considered differentially expressed proteins (DEPs); no fold-change threshold was applied. Positive log_2_ fold changes indicate higher abundance in females (sex comparisons) or in the DREADD group (genotype comparisons). Results of each comparison were displayed as volcano plots on common axes, indicating nominal P = 0.05 and the P-value corresponding to FDR = 0.05 (the largest P-value among proteins with an adjusted P < 0.05 or, when no protein reached this threshold, 0.05 divided by the number of proteins tested). For each comparison, a PCA of the animals of the two compared groups was calculated (prcomp; mean-centred, unscaled log_2_ intensities of the proteins quantified in all of these animals).

#### Software and data availability

Analyses were performed in R with the packages limma, readxl, openxlsx, ggplot2 and ggrepel; MaxLFQ intensities were calculated with the diann R package. The MaxLFQ intensity tables and all analysis results are provided in **Supplementary Data 1**. The complete R code is provided in **Supplementary Data 2**. The analysis code was developed with the assistance of an AI model (Claude, Anthropic), which was used to write, debug and document the R scripts. The mass spectrometry proteomics data have been deposited to the ProteomeXchange Consortium via the PRIDE (*51*) partner repository with the dataset identifier PXD080330.

### Assessment of mechanical withdrawal thresholds

For behavioral testing, male and female rats were habituated to both the testing environment and the experimenter on two consecutive days. Baseline mechanical withdrawal thresholds were then determined using the modified up-and-down von Frey method (*52*) on two subsequent days (Baseline 1 and Baseline 2). Animals subsequently received spinal intraparenchymal injections of either Gq-DREADD, Gi-DREADD, or control viral vectors. Ten days later, mechanical thresholds were reassessed immediately prior to intraperitoneal administration of CNO (3 mg kg^-^¹; Baseline 3). Mechanical threshold measurements were then performed 1 -2 h, 4 -5 h, 24 h, and 48 h after CNO administration.

### Immunostainings

For immunostaining, animals were deeply anaesthetized with isoflurane (5 vol%), followed by intraperitoneal injection of ketamine (50 mgꞏkg^-^¹) and xylazine (5 mgꞏkg^-^¹), and transcardially perfused with ice-cold 0.9% NaCl containing 1% heparin, followed by 4% paraformaldehyde (PFA, pH 8.4). Spinal cords were dissected and post-fixed overnight in PFA at 4 °C, then cryoprotected sequentially in 20% and 30% sucrose in 0.1 M phosphate-buffered saline (PBS) for 24 h each. Tissue was subsequently embedded in optimal cutting temperature compound Cryostat Embedding Medium (OCT, Scigen Scientific Gardena, 23-730-625), flash-frozen in isopentane, and stored at −80 °C until sectioning. Transverse sections (40 µm) were cut using a cryostat and stored in well plates containing 0.05% sodium azide in PBS at 4 °C.

Free-floating indirect immunohistochemical staining was performed on collected tissue sections. Sections were washed in PBS containing 0.1% Triton X-100 (3 × 10 minutes) and subsequently incubated for 60 minutes in blocking solution consisting of 4% normal goat serum (Cell Signaling Technology, 5425) in PBS containing 0.1% Triton X-100. All washing and incubation steps were performed at room temperature on a shaker (70 rpm).

Sections were then incubated overnight with primary antibodies and/or biotinylated Wisteria floribunda agglutinin diluted in blocking solution: aggrecan (Invitrogen, MA3-16888, mouse monoclonal, 1:75), androgen receptor (Invitrogen, PA1-110, rabbit polyclonal, 1:150), GFAP (Thermo Fisher Scientific, PA1-10004, chicken polyclonal, 1:1000), GFAP (Cell Signaling, 12389, rabbit monoclonal, 1:500), NeuN (Millipore, MAB377, mouse monoclonal, 1:600), NeuN (Cell Signaling, 12943S, rabbit monoclonal, 1:1000), NK1R (Millipore, AB15810, guinea pig polyclonal, 1:1000), S100β (Synaptic Systems, 287004, guinea pig monoclonal, 1:500), and biotinylated Wisteria floribunda agglutinin (Vector Laboratories, B-1355-2, 1:1000).

Following incubation with primary antibodies and/or biotinylated Wisteria floribunda agglutinin, sections were washed in PBS containing 0.1% Triton X-100 (3 × 10 minutes) and incubated for 2 hours at room temperature with the appropriate fluorophore-conjugated secondary antibodies or streptavidin diluted in blocking solution: Alexa Fluor 488 goat anti-guinea pig (Thermo Fisher Scientific, A-11073, 1:500), Alexa Fluor 488 goat anti-chicken (Thermo Fisher Scientific, A-11039, 1:500), Alexa Fluor 488 streptavidin (Thermo Fisher Scientific, S32354, 1:250), Alexa Fluor 488 goat anti-rabbit (Thermo Fisher Scientific, A-11008, 1:1000), Alexa Fluor 546 goat anti-mouse (Thermo Fisher Scientific, A-11003, 1:500), Alexa Fluor 647 goat anti-guinea pig (Thermo Fisher Scientific, A-21450, 1:250), Alexa Fluor 647 goat anti-rabbit (Thermo Fisher Scientific, A-21245, 1:500), Alexa Fluor 647 goat anti-mouse (Invitrogen, A-21235, 1:1000), Brilliant Violet 421 donkey anti-rabbit (BioLegend, 406410, 1:100), and Brilliant Violet 421 streptavidin (BioLegend, 405226, 1:1000). All antibodies and reagents used are also summarized in **Supplementary Table 10**.

Sections were subsequently washed in PBS (3 × 10 minutes) and incubated with 0.2% DAPI (Abcam, ab228549) in PBS for 60 minutes. For stainings involving Brilliant Violet fluorophores, DAPI staining was omitted. For neuronal network stainings, Triton X-100 was omitted from all steps of the protocol.

Finally, sections were washed in PBS (2 × 5 minutes), mounted onto glass slides, coverslipped using Fluoromount-G mounting medium (Invitrogen, 4958-02), and stored at 4 °C until imaging. Confocal z-stacks were acquired at 1 μm intervals using a Zeiss LSM 780 inverted confocal microscope. High-resolution three-dimensional image reconstructions were generated and processed using ZEN software (version 2.3 SP1). All images were analyzed using IMARIS x64 software (version 10.1.0, Oxford Instruments). For analysis of viral specificity, NeuN-positive cells and mCherry-positive virus-transduced cells were identified using the Spots tool in Imaris. Colocalization was determined using the ‘shortest distance to Spots’ filter, with a maximum distance threshold of 6 µm, followed by manual verification. For analysis of androgen receptor (AR) expression, NeuN-positive neurons and astrocytes co-expressing GFAP and S100β were identified using the Spots tool in Imaris and assessed for AR immunoreactive signal. Colocalization with AR signal was determined using the same ‘shortest distance to Spots’ approach, with a maximum distance threshold of 5 µm, followed by manual verification. For quantification of perineuronal net (PNN) intensity surrounding CTB-positive projection neurons, individual neurons were first reconstructed by generating automatic surface objects in the CTB channel. If more than two neurons were merged into a single surface object within one tissue section, individual neuronal surfaces were separated manually using the ‘Split Surface’ function. To assess the extracellular matrix surrounding each neuron, the ‘Dilate Surface’ extension (Open Source Extensions, Oxford Instruments) was used to generate a perisomatic ring around each neuronal surface at a user-defined distance. Mean fluorescence intensity within the resulting dilated surface was then extracted from the corresponding WFA or aggrecan channel and used to quantify PNN labeling around CTB-positive projection neurons. For quantification of PNN intensity in the whole SDH, regions of interest were manually defined using surface objects based on all available channels to delineate the superficial dorsal horn. Mean fluorescence intensity within these regions was extracted from the corresponding WFA and aggrecan channels and used for quantification of PNNs in laminae I–II.

### Cytokine Assay

Ten to twelve days after spinal viral injections, spinal cords were rapidly removed and 500 µm transverse sections were prepared as described above. Sections exhibiting mCherry fluorescence were identified, and the ipsilateral dorsal horn was microdissected under a stereomicroscope. As a control, contralateral dorsal horn sections were also microdissected. For cytokine measurements, six sections from ipsi-and contralateral side from each animal were transferred into two wells of a 48-well plate containing 150 µl HEPES-buffered solution (in mM: 145 NaCl, 1.9 KCl, 2.4 CaCl₂, 1.3 MgCl₂, 10 HEPES, and 10 glucose; pH 7.38; 310-320 mOsmolꞏkg⁻¹). Following application of CNO (10 µM) in HEPES or HEPES alone, sections were incubated for 2 hours at room temperature. The supernatant was then collected, immediately frozen, and stored at −80 °C until analysis. Cytokine concentrations were quantified using the Bio-Plex Pro™ Rat Cytokine 23-Plex Assay (Bio-Rad, #12005641) according to the manufacturer’s instructions.

### Image Creation

Animal schematics and selected graphical elements were manually drawn in CorelDRAW® Graphics Suite 2019, which was also used for final figure assembly (version 21.0.0.593).

## Acknowledgments

We thank Nicole Kanta and Elisabeth Barnreiter (University of Vienna) for excellent technical assistance, Barbara Jilka for help with image analysis, and Roni Hogri for support with the analysis of calcium imaging data, and Igor Adameyko for granting access to the confocal microscope. We are grateful to the staff of the animal facility at the Center for Brain Research for expert animal care and support. We thank all members of the Drdla-Schutting laboratory for helpful discussions. The proteomic results presented have been achieved in part using the Austrian Scientific Cluster (ASC).

## Author contributions

Conceptualization and funding acquisition: RDS, MS

Electrophysiology in vitro: LK, SA

Electrophysiology in vivo: RDS

Calcium imaging: RDS, RH, ER

Behavioral experiments: DK, LK

Immunostainings, microscopy: ASMS, LT, ER, LK

Image analysis: ASMS, LT, DK, RH, LK, ER

Intraspinal injections: RH, ASMS, SA, LK

Supraspinal injections: RH, LK

Proteomics: LU, MS

Cytokine Assays: PWS

Supervision: RDS

Writing -original draft: RDS, MS

Writing -review and editing: RDS, MS, LK, LU, with input from all authors

## Data, code, and materials availability

The MS proteomics raw data, including metadata, and protein matrices for each dataset are available via ProteomeXchange with identifier PXD080330 (also see Supplementary Data 1 and 2). All other data supporting the findings of this study are included in the manuscript (and Supplementary Table 1-9), raw data are available from the corresponding author upon reasonable request.

## Supplementary Information

Extended Data Figures 1 to 6

Supplementary Tables 1 to 10

Supplementary Data 1 to 2

## References and Notes

1. R.-R. Ji, T. Berta, M. Nedergaard, Glia and pain: Is chronic pain a gliopathy? PAIN® 154, S10–S28 (2013).

2. J. S. Mogil, M. Parisien, S. J. Esfahani, L. Diatchenko, Sex differences in mechanisms of pain hypersensitivity. Neuroscience & Biobehavioral Reviews 163, 105749 (2024).

3. M. Bortolanza, D. Gobbo, L. Fang, A. Scheller, X. Bai, F. Kirchhoff, Sex-Specific Properties of Astrocytes: From Development to Evolutionary Insights. Neurochem Res 50, 267 (2025).

4. E. Gozlan, Y. Lewit-Cohen, D. Frenkel, Sex Differences in Astrocyte Activity. Cells 13, 1724 (2024).

5. A. Y. Zhang, E. Elias, M. T. Manners, Sex-dependent astrocyte reactivity: Unveiling chronic stress-induced morphological changes across multiple brain regions. Neurobiology of Disease 200, 106610 (2024).

6. M. M. Halassa, T. Fellin, P. G. Haydon, The tripartite synapse: roles for gliotransmission in health and disease. Trends in Molecular Medicine 13, 54–63 (2007).

7. G. Perea, M. Navarrete, A. Araque, Tripartite synapses: astrocytes process and control synaptic information. Trends in Neurosciences 32, 421–431 (2009).

8. P. Kofuji, A. Araque, G-Protein-Coupled Receptors in Astrocyte-Neuron Communication. Neuroscience 456, 71–84 (2021).

9. A. Verkhratsky, M. Nedergaard, Physiology of Astroglia. Physiol Rev 98, 239– 389 (2018).

10. E. Dzyubenko, D. M. Hermann, Role of glia and extracellular matrix in controlling neuroplasticity in the central nervous system. Semin Immunopathol 45, 377– 387 (2023).

11. S. Ada, L. Klinger, H. L. Teuchmann, V. Mussetto, V. Hadschieff, M. T. Kronschläger, A. S. M. Siegert, L. Trofimova, R. Holzinger, D. Kurija, R. Drdla-Schutting, Astrocytogenic bidirectional plasticity at spinal nociceptive synapses regulates acute nociceptive processing. PAIN 167, 372 (2026).

12. A. M. Barry, N. Zhao, X. Yang, D. L. Bennett, G. Baskozos, Deep RNA-seq of male and female murine sensory neuron subtypes after nerve injury. Pain 164, 2196– 2215 (2023).

13. J. R. Sondermann, A. M. Barry, F. Xian, T. Haberl, D. Gomez-Varela, F. Benseler, D. Schreiner, N. Brose, N. Lipstein, M. Schmidt, Proximity labelling reveals the compartmental proteome of murine sensory neurons. bioRxiv [Preprint] (2025). 10.1101/2025.06.13.659475.

14. J. Jabłońska, G. Wiera, J. W. Mozrzymas, Extracellular matrix integrity regulates GABAergic plasticity in the hippocampus. Matrix Biology 134, 184–196 (2024).

15. K. M. Kamal, A. R. Ghazali, G. T. Selvarajah, N. S. A. Mutalib, N. Abu, E. W. Chua, S. F. Masre, The extracellular matrix: structure, composition, biological functions, diseases, and therapeutic targets. Mol Biomed 7, 38 (2026).

16. J. W. Fawcett, M. Fyhn, P. Jendelova, J. C. F. Kwok, J. Ruzicka, B. A. Sorg, The extracellular matrix and perineuronal nets in memory. Mol Psychiatry 27, 3192–3203 (2022).

17. B. P. Tewari, A. M. Woo, C. E. Prim, L. Chaunsali, D. C. Patel, I. F. Kimbrough, K. Engel, J. L. Browning, S. L. Campbell, H. Sontheimer, Astrocytes require perineuronal nets to maintain synaptic homeostasis in mice. Nat Neurosci 27, 1475–1488 (2024).

18. J. Sánchez-Ventura, M. A. Lane, E. Udina, The Role and Modulation of Spinal Perineuronal Nets in the Healthy and Injured Spinal Cord. Front Cell Neurosci 16, 893857 (2022).

19. S. Tansley, N. Gu, A. U. Guzmán, W. Cai, C. Wong, K. C. Lister, E. Muñoz-Pino, N. Yousefpour, R. B. Roome, J. Heal, N. Wu, A. Castonguay, G. Lean, E. M. Muir, A. Kania, M. Prager-Khoutorsky, J. Zhang, C. G. Gkogkas, J. W. Fawcett, L. Diatchenko, A. Ribeiro-da-Silva, Y. De Koninck, J. S. Mogil, A. Khoutorsky, Microglia-mediated degradation of perineuronal nets promotes pain. Science 377, 80–86 (2022).

20. G. Cornez, O. T. Shevchouk, S. Ghorbanpoor, G. F. Ball, C. A. Cornil, J. Balthazart, Testosterone stimulates perineuronal nets development around parvalbumin cells in the adult canary brain in parallel with song crystallization. Hormones and Behavior 119, 104643 (2020).

21. S. Soleman, P. K. Yip, D. A. Duricki, L. D. F. Moon, Delayed treatment with Chondroitinase ABC promotes sensorimotor recovery and plasticity after stroke in aged rats. Brain 135, 1210–1223 (2012).

22. J. M. Massey, C. H. Hubscher, M. R. Wagoner, J. A. Decker, J. Amps, J. Silver, S. M. Onifer, Chondroitinase ABC Digestion of the Perineuronal Net Promotes Functional Collateral Sprouting in the Cuneate Nucleus after Cervical Spinal Cord Injury. J. Neurosci. 26, 4406–4414 (2006).

23. P. Chu, R. Abraham, K. Budhu, U. Khan, N. De Marco Garcia, J. C. Brumberg, The Impact of Perineuronal Net Digestion Using Chondroitinase ABC on the Intrinsic Physiology of Cortical Neurons. Neuroscience 388, 23–35 (2018).

24. T. V. Vaidyanathan, M. Collard, S. Yokoyama, M. E. Reitman, K. E. Poskanzer, Cortical astrocytes independently regulate sleep depth and duration via separate GPCR pathways. Elife 10, e63329 (2021).

25. A. Covelo, A. Araque, Neuronal activity determines distinct gliotransmitter release from a single astrocyte. eLife 7, e32237 (2018).

26. D. N. Xanthos, J. Sandkühler, Neurogenic neuroinflammation: inflammatory CNS reactions in response to neuronal activity. Nat Rev Neurosci 15, 43–53 (2014).

27. N. Zhang, Z. Yan, H. Liu, M. Yu, Y. He, H. Liu, C. Liang, L. Tu, L. Wang, N. Yin, J. Han, N. Scarcelli, Y. Yang, C. Wang, T. Zeng, L.-L. Chen, Y. Xu, Hypothalamic Perineuronal Nets Are Regulated by Sex and Dietary Interventions. Front Physiol 12, 714104 (2021).

28. N. Kiguchi, M. Miyajima, F. Saika, Y. Fukazawa, S. Hino, K. Suzuki, Expression of androgen receptors in the spinal dorsal horn neurons of mice. Pain Research 40, 135– 138 (2025).

29. E. Dzyubenko, D. M. Hermann, Role of glia and extracellular matrix in controlling neuroplasticity in the central nervous system. Semin Immunopathol 45, 377– 387 (2023).

30. O. Bukalo, M. Schachner, A. Dityatev, Modification of extracellular matrix by enzymatic removal of chondroitin sulfate and by lack of tenascin-R differentially affects several forms of synaptic plasticity in the hippocampus. Neuroscience 104, 359–369 (2001).

31. J. Jabłońska, G. Wiera, J. W. Mozrzymas, Extracellular matrix integrity regulates GABAergic plasticity in the hippocampus. Matrix Biol 134, 184–196 (2024).

32. P. A. McRae, B. E. Porter, The Perineuronal Net Component of the Extracellular Matrix in Plasticity and Epilepsy. Neurochem Int 61, 963–972 (2012).

33. J. Lu, L. Yang, Y. Xu, L. Ai, J. Chen, F. Xiong, L. Hu, H. Chen, J. Liu, X. Yan, H. Huang, L. Chen, C. Yu, The Modulatory Effect of Motor Cortex Astrocytes on Diabetic Neuropathic Pain. J. Neurosci. 41, 5287–5302 (2021).

34. Q. Xu, N. C. Ford, S. He, Q. Huang, M. Anderson, Z. Chen, F. Yang, L. K. Crawford, M. J. Caterina, Y. Guan, X. Dong, Astrocytes contribute to pain gating in the spinal cord. Sci Adv 7, eabi6287 (2021).

35. M. J. Pereira, R. Ayana, M. G. Holt, L. Arckens, Chemogenetic manipulation of astrocyte activity at the synapse— a gateway to manage brain disease. Front. Cell Dev. Biol. 11 (2023).

36. C. A. Durkee, A. Covelo, J. Lines, P. Kofuji, J. Aguilar, A. Araque, Gi/o protein-coupled receptors inhibit neurons but activate astrocytes and stimulate gliotransmission. Glia 67, 1076–1093 (2019).

37. T. Gould, L. Chen, Z. Emri, T. Pirttimaki, A. C. Errington, V. Crunelli, H. R. Parri, GABAB receptor-mediated activation of astrocytes by gamma-hydroxybutyric acid. Philosophical Transactions of the Royal Society B: Biological Sciences 369, 20130607 (2014).

38. L. Mariotti, G. Losi, M. Sessolo, I. Marcon, G. Carmignoto, The inhibitory neurotransmitter GABA evokes long-lasting Ca(2+) oscillations in cortical astrocytes. Glia 64, 363–373 (2016).

39. I. Savtchouk, A. Volterra, Gliotransmission: Beyond Black-and-White. J Neurosci 38, 14–25 (2018).

40. A. Serrano, N. Haddjeri, J.-C. Lacaille, R. Robitaille, GABAergic Network Activation of Glial Cells Underlies Hippocampal Heterosynaptic Depression. J. Neurosci. 26, 5370–5382 (2006).

41. S. N. Heimbach, Á. C. Matute, V. Steininger, R. Rajadhyaksha, L. Klein, L. Ferguson, Y. A. Sabir, M. Huang, A. Cruz-Martín, S. Melzer, Temporal cortex astrocytic Gi-GPCR signaling regulates learned threat responses. bioRxiv [Preprint] (2026). 10.64898/2026.05.21.724789.

42. Y. Van Den Herrewegen, T. M. Sanderson, S. Sahu, D. De Bundel, Z. A. Bortolotto, I. Smolders, Side-by-side comparison of the effects of Gq-and Gi-DREADD-mediated astrocyte modulation on intracellular calcium dynamics and synaptic plasticity in the hippocampal CA1. Molecular Brain 14, 144 (2021).

43. R. Hogri, H. L. Teuchmann, B. Heinke, R. Holzinger, L. Trofimova, J. Sandkühler, GABAergic CaMKIIα+ Amygdala Output Attenuates Pain and Modulates Emotional-Motivational Behavior via Parabrachial Inhibition. J Neurosci 42, 5373–5388 (2022).

44. G. Paxinos, C. Watson, The Rat Brain in Stereotaxic Coordinates (Elsevier/Academic, 2009; https://books.google.at/books?id=sQNfmwEACAAJ).

45. C. S. Hughes, S. Moggridge, T. Müller, P. H. Sorensen, G. B. Morin, J. Krijgsveld, Single-pot, solid-phase-enhanced sample preparation for proteomics experiments. Nat Protoc 14, 68–85 (2019).

46. V. Demichev, C. B. Messner, S. I. Vernardis, K. S. Lilley, M. Ralser, DIA-NN: neural networks and interference correction enable deep proteome coverage in high throughput. Nat Methods 17, 41–44 (2020).

47. F. Kistner, J. L. Grossmann, L. R. Sinn, V. Demichev, QuantUMS: uncertainty minimisation enables confident quantification in proteomics. bioRxiv [Preprint] (2023). 10.1101/2023.06.20.545604.

48. J. Cox, M. Y. Hein, C. A. Luber, I. Paron, N. Nagaraj, M. Mann, Accurate Proteome-wide Label-free Quantification by Delayed Normalization and Maximal Peptide Ratio Extraction, Termed MaxLFQ*. Molecular & Cellular Proteomics 13, 2513–2526 (2014).

49. R Core Team (2024) R A Language and Environment for Statistical Computing. R Foundation for Statistical Computing. -References -Scientific Research Publishing. https://www.scirp.org/reference/referencespapers?referenceid=3887298.

50. M. E. Ritchie, B. Phipson, D. Wu, Y. Hu, C. W. Law, W. Shi, G. K. Smyth, limma powers differential expression analyses for RNA-sequencing and microarray studies. Nucleic Acids Res 43, e47 (2015).

51. Y. Perez-Riverol, C. Bandla, D. J. Kundu, S. Kamatchinathan, J. Bai, S. Hewapathirana, N. S. John, A. Prakash, M. Walzer, S. Wang, J. A. Vizcaíno, The PRIDE database at 20 years: 2025 update. Nucleic Acids Res 53, D543–D553 (2025).

52. R. P. Bonin, C. Bories, Y. De Koninck, A simplified up-down method (SUDO) for measuring mechanical nociception in rodents using von Frey filaments. Mol Pain 10, 26 (2014).

