## Extended Data for "Sex-dependent gating of astrocyte-driven nociceptive plasticity"

Laura Klinger *et al.*

**This PDF file includes:**

Extended Data Figures 1 to 6

**Other Supplementary Information for this manuscript include the following:**

Supplementary Table 1 to 10
Supplementary Data 1 to 2

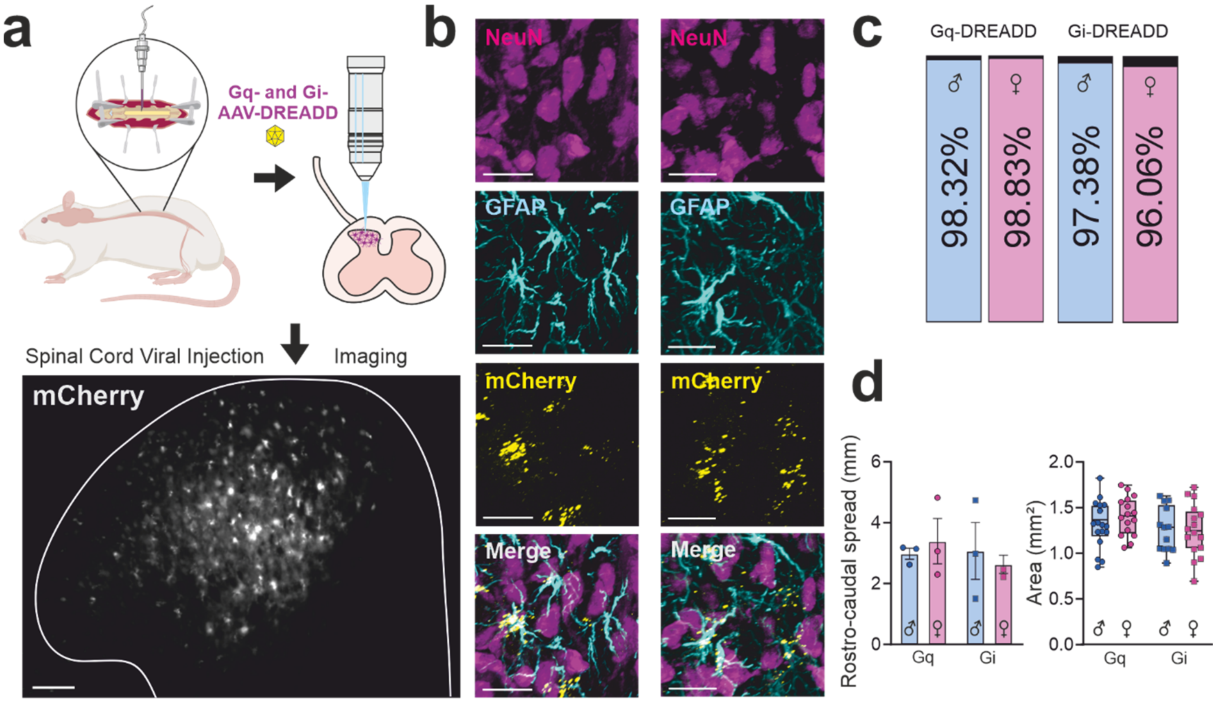


**Extended Data Fig. 1. DREADDs are selectively targeted to spinal dorsal horn astrocytes.** (**a**) DREADDs were delivered to the spinal dorsal horn by intraparenchymal AAV injection. A representative fluorescence image illustrates mCherry expression in the dorsal horn. Scale bar, 100 µm. (**b**) Representative confocal images showing immunostaining for the neuronal marker NeuN (magenta) and the astrocytic marker GFAP (cyan) together with DREADD expression (mCherry, yellow) in Gi-DREADD (left) and Gq-DREADD (right) animals. Scale bars, 20 µm. (**c**) Quantification of NeuN/mCherry co-localization demonstrates minimal neuronal expression of the viral constructs. Values indicate the percentage of mCherry-positive cells that were NeuN-negative. (**d**) Left: Quantification of the rostrocaudal spread of DREADD expression in Gq-DREADD (3 males, 3 females) and Gi-DREADD (3 males, 3 females) animals. Data are shown as mean ± SEM. Right: Quantification of the area of DREADD expression within the dorsal horn, defined as the mCherry-positive area, in Gq-DREADD (males: n = 13 slices from 4 animals; females: n = 16 slices from 4 animals) and Gi-DREADD (males: n = 17 slices from 5 animals; females: n = 16 slices from 4 animals) animals. Details are given in **Supplementary Table 6**.

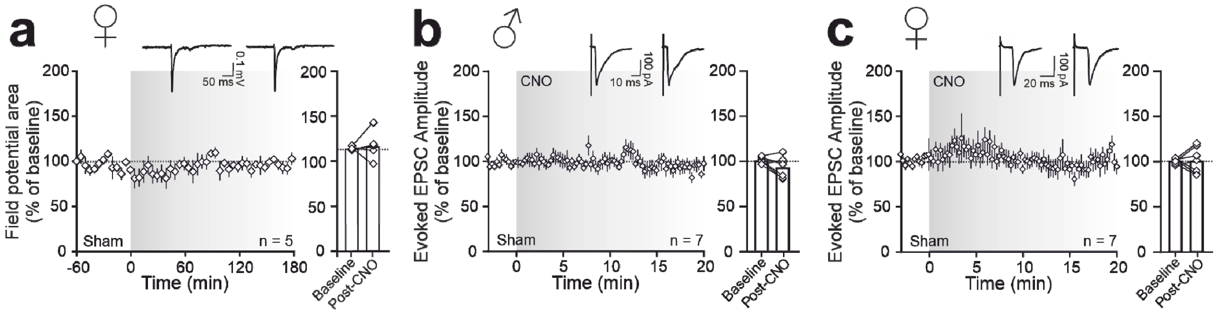


**Extended Data Fig. 2. Sham virus controls show no effect of CNO on synaptic transmission.** (**a**) Time course of C-fiber-evoked field potentials recorded in vivo from the SDH of female rats expressing a sham virus lacking DREADDs. Systemic administration of CNO (grey shading) did not alter field potential area. Representative traces recorded before and after CNO application are shown above the graph. The bar graph summarizes the mean field potential area during the baseline and post-CNO periods. (**b**, **c**) Time course of monosynaptic C-fiber-evoked EPSC amplitudes recorded from superficial dorsal horn neurons in acute spinal cord slices obtained from male (**b**) and female (**c**) sham-virus animals. Bath application of CNO (10 µM; grey shading) did not affect EPSC amplitudes in either sex. Representative EPSC traces recorded before and after CNO application are shown above each graph. Bar graphs summarize mean EPSC amplitudes during baseline and after CNO application. Data are normalized to baseline and presented as mean ± SEM. Individual data points in bar graphs represent single experiments. CNO had no significant effect on synaptic transmission in sham-virus control animals, confirming that the synaptic changes observed in DREADD-expressing animals were dependent on chemogenetic receptor activation. Details are given in **Supplementary Table 7**.

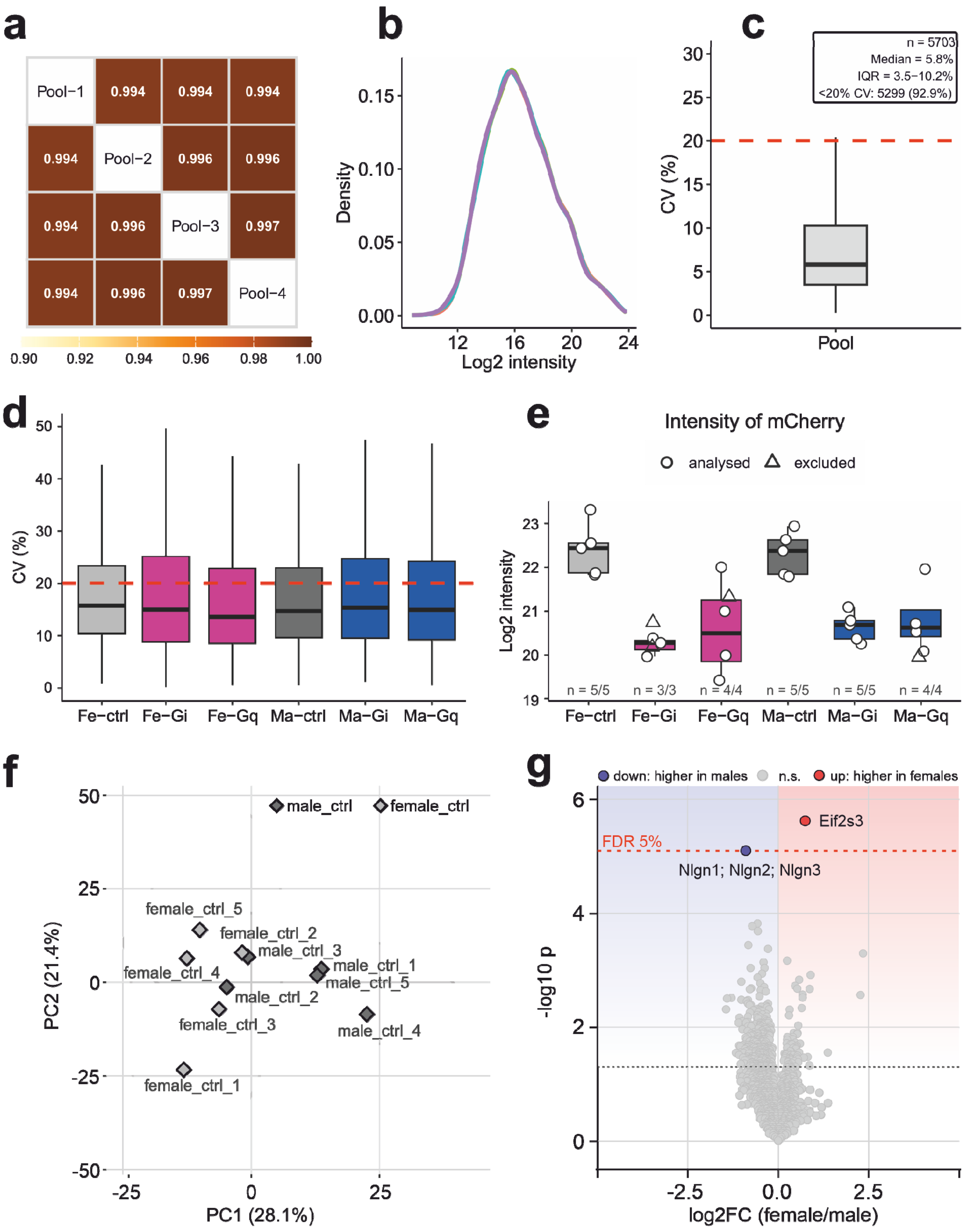


**Extended Data Fig. 3. Technical quality control of the proteomic dataset and comparison of female and male control animals.** (**a**) Pearson correlation coefficients of $log_2$-transformed MaxLFQ intensities across four repeated injections of a pooled quality-control (QC) sample (Pool-1 to Pool-4). (**b**) Density distributions of $log_2$-transformed intensities for each QC injection. (**c**) Coefficients of variation (CVs) across QC injections for gene groups quantified in at least three of the four injections. The inset indicates the number of gene groups, median CV, interquartile range (IQR), and number and percentage of gene groups with a CV below 20%. (**d**) CVs within each sex × genotype group, calculated for gene groups quantified in at least three analyzed animals per group. (**e**) $Log_2$-transformed intensity of the mCherry reporter (PAmCherry) in individual animals. Circles indicate animals included in the final analysis; triangles indicate samples excluded due to high blood content. Numbers indicate the total number of quantified animals and the number included in the final analysis for each group. (**f**) Principal component analysis (PCA) of male and female control samples based on mean-centered $log_2$-transformed intensities of 4987 gene groups quantified in all 10 animals. Ellipses indicate 95% confidence regions for each group, and percentages indicate the variance explained by each principal component. (**g**) Volcano plot comparing female and male control samples (5283 gene groups tested). Positive $log_2$ fold changes ($log_2FC$) indicate higher abundance in females. Horizontal dotted lines indicate the P-value thresholds corresponding to a false discovery rate (FDR) of 0.05 (upper) and nominal $P = 0.05$ (lower). Gene groups with $FDR < 0.05$ are highlighted in red (higher abundance in females) or blue (higher abundance in males). Further details are provided in **Supplementary Data 1 and 2**.

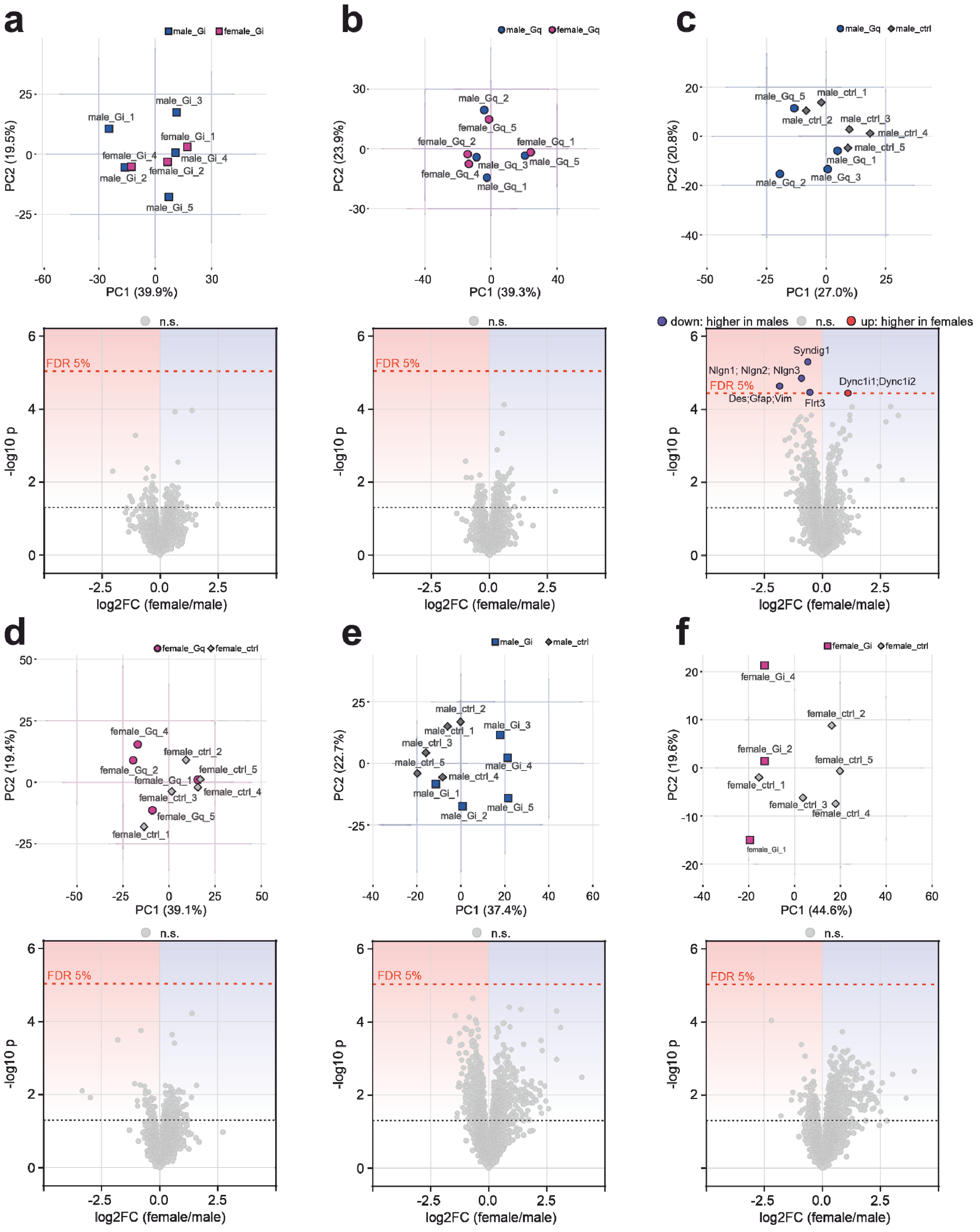


**Extended Data Fig. 4. Female versus male comparisons in DREADD animals and DREADD versus control comparisons within each sex.** (**a**) Female versus male Gi animals (female n = 3, male n = 5; PCA, 5199 gene groups; volcano plot, 5371 gene groups tested). (**b**) Female versus male Gq animals (n = 4 each; 5095 and 5426 gene groups). (**c**) Gq versus control males (Gq n = 4, control n = 5; 5046 and 5338 gene groups). (**d**) Gq versus control females (Gq n = 4, control n = 5; 5029 and 5351 gene groups). (**e**) Gi versus control males (n = 5 each; 5046 and 5318 gene groups). (**f**) Gi versus control females (Gi n = 3, control n = 5; 5057 and 5268 gene groups). In each panel, the top plot shows a principal component analysis (PCA) of the animals of the two compared groups, based on the mean-centered $\log_2$ intensities of the gene groups quantified in all of these animals; ellipses show 95% confidence regions for groups with more than three animals, and percentages give the variance explained by each component. The bottom plot shows the corresponding volcano plot; positive $\log_2$ fold change ($\log_2$FC) indicates higher abundance in females (**a** and **b**) or in the DREADD group (**c** to **f**). Horizontal dotted lines mark the P value corresponding to a false discovery rate (FDR) of 0.05 (upper) and nominal P = 0.05 (lower). Proteins with FDR < 0.05 are colored (red, higher; blue, lower) and labelled. Details are given in **Supplementary Data 1 and 2**.

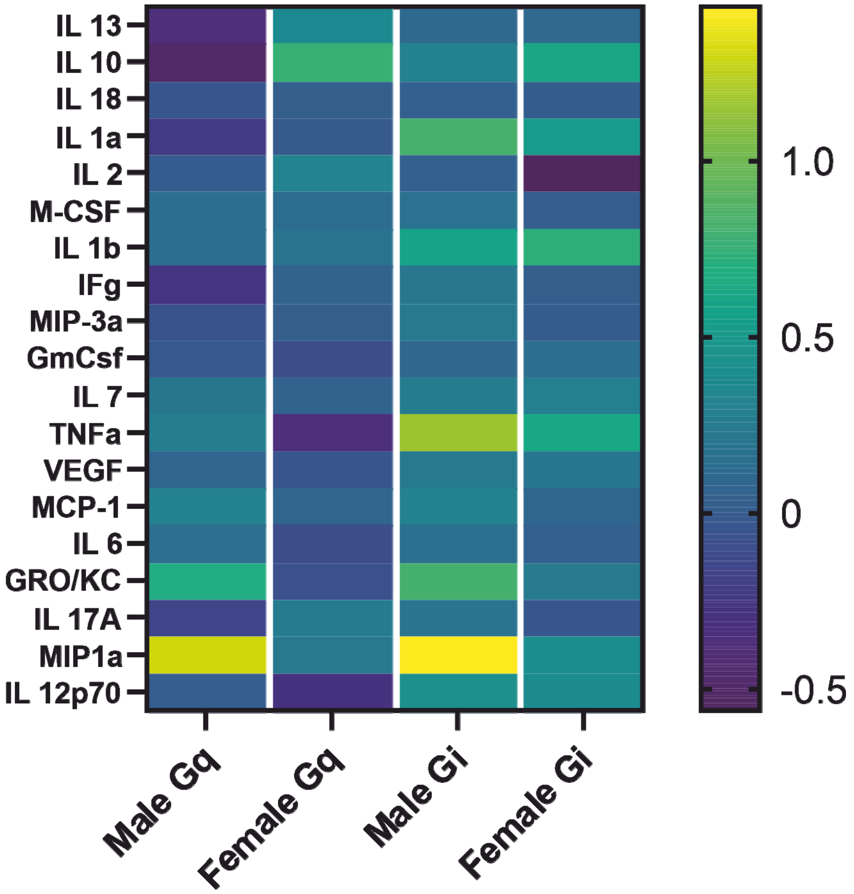


**Extended Data Fig. 5. Cytokine profiles following astrocytic Gi- and Gq-DREADD activation in male and female rats.** Heatmaps show $log_2$-transformed mean fold changes in cytokine and chemokine concentrations measured in whole spinal dorsal horn slice supernatants after CNO application. Technical duplicates were averaged, and individual fold changes were calculated relative to the corresponding same-sex control animal before averaging within each group (n = 2 DREADD animals per sex and condition). Positive and negative values indicate higher and lower concentrations relative to controls, respectively; zero indicates no change. Analytes with insufficient quantifiable data were omitted from the heatmaps. Comparisons between males and females were performed separately within each DREADD condition using unpaired Welch's t-tests on individual, untransformed fold changes. No statistically significant sex differences were detected. Individual values and unadjusted p-values are provided in **Supplementary Table 8**.

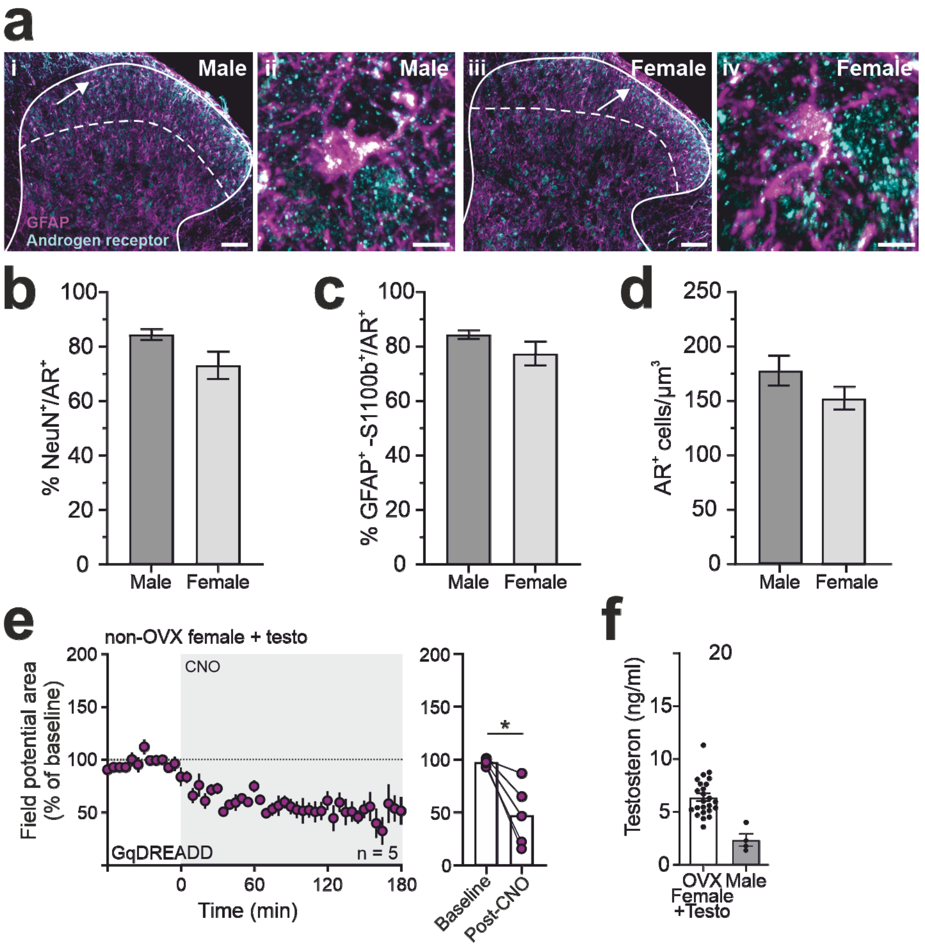


**Extended Data Fig. 6. Androgen receptor expression in the spinal dorsal horn and testosterone-dependent astrocyte-driven synaptic plasticity.** (**a**) Representative confocal images of lumbar spinal dorsal horn sections from male (i, ii) and female (iii, iv) rats stained for GFAP (magenta) and androgen receptor (AR; cyan). Low-magnification images (i, iii) illustrate AR expression throughout the superficial dorsal horn (outlined; the dashed line indicates the approximate border between superficial and deeper laminae), whereas high-magnification images (ii, iv) show AR-positive cells relative to GFAP-positive astrocytes. Scale bars: 100 µm (i, iii), 10 µm (ii, iv). (**b**) Quantification of the percentage of NeuN-positive neurons expressing AR in the superficial dorsal horn of male and female rats. (**c**) Quantification of the percentage of GFAP- and S100β-positive astrocytes expressing AR. (**d**) Density of AR-positive cells in the superficial dorsal horn. No significant sex differences were detected in neuronal or astrocytic AR expression or in the overall density of AR-positive cells. (**e**) Time course of C-fiber-evoked field potential area recorded in the spinal dorsal horn of testosterone-treated, non-ovariectomized (OVX) female rats expressing astrocytic Gq-DREADDs. Systemic CNO application (3 mg·kg·h-1, i.v.), grey area, induced a sustained depression of C-fiber-evoked field potentials. Data are expressed as percentages of baseline (mean ± SEM; n = 5 animals). Right, individual values before (Baseline) and after CNO application (Post-CNO). Statistics: paired T-Test; $*P < 0.05$. (**f**) Serum testosterone concentrations in OVX female rats treated with testosterone and untreated male rats. Individual data points represent single animals; bars indicate mean ± SEM. Details are given in **Supplementary Table 9**.
