## Supplementary Information for "Sex-dependent gating of astrocyte-driven nociceptive plasticity"

Laura Klinger *et al.*

**This PDF file includes:**

Supplementary Table 1 to 10

**Other Supplementary Information for this manuscript include the following:**

Extended Data Figures 1 to 6
Supplementary Data 1 to 2

**Supplementary Table 1. Electrophysiological recordings in vivo (paired t-tests).**

| Figure | Treatment | Mean ± SEM (%) | N (animals) | P-Value (paired t-test) | t, df |
|---|---|---|---|---|---|
| Fig. 1b | Gq-DREADD/CNO Male | Baseline: 99 ± 3<br>Post-CNO: 68 ± 7 | 7 | 0.0007 | t=6.417, df=6 |
| Fig. 1c | Gq-DREADD/CNO Female | Baseline: 100 ± 1<br>Post-CNO: 96 ± 11 | 7 | 0.6854 | t=0.4254, df=6 |
| Fig. 1d | Gi-DREADD/CNO Male | Baseline: 101 ± 1<br>Post-CNO: 174 ± 23 | 7 | 0.0185 | t=3.206, df=6 |
| Fig. 1e | Gi-DREADD/CNO Female | Baseline: 100 ± 2<br>Post-CNO: 112 ± 9 | 8 | 0.2575 | t=1.233, df=7 |

**Supplementary Table 2. Membrane Potential, Cell Capacitance, Paired pulse ratio and evoked EPSCs of recorded cells.**

| Figure | Treatment | N (cells) | Resting Membrane Potential (mV) | Cell Capacitance (pF) | Paired Puls ratio (paired t-test) Mean ± SEM (%) | p-Value (paired t-test) | t, df | evoked EPSCs (paired t-test) Mean ± SEM (%) | p-Value (paired t-test) | t, df |
|---|---|---|---|---|---|---|---|---|---|---|
| Fig. 2b | C-fiber DREADD Gq male | 8 | -47 ± 3.27 | 42 ± 5.3 | Baseline: 1.9 ± 0.6 Post-CNO: 1.9 ± 0.6 | 0.7274 | t=0.3628, df=7 | Baseline: 99 ± 1.8 Post-CNO: 77 ± 2.8 | 0.0001 | t=7.525, df=7 |
| Fig. 2c | C-fiber DREADD Gq female | 7 | -47 ± 1.01 | 46 ± 5.1 | Baseline: 0.8 ± 0.08 Post-CNO: 0.8 ± 0.06 | 0.7288 | t=0.3634, df=6 | Baseline: 100 ± 1.9 Post-CNO: 95 ± 5 | 0.4509 | t=0.8062, df=6 |
| Fig. 2d | C-fiber DREADD Gi male | 9 | -48 ± 2.05 | 57 ± 8.4 | Baseline: 0.74 ± 0.1 Post-CNO: 0.55 ± 0.05 | 0.0947 | t=1.895, df=8 | Baseline: 101 ± 1.3 Post-CNO: 133 ± 8 | 0.0042 | t=3.960, df=8 |
| Fig. 2e | C-fiber DREADD Gi female | 12 | -47 ± 1.5 | 67 ± 6.2 | Baseline: 1.4 ± 0.6 Post-CNO: 1.3 ± 0.5 | 0.0542 | t=2.155, df=11 | Baseline: 102 ± 1.2 Post-CNO: 99 ± 3.9 | 0.5522 | t=0.6132, df=11 |
| Fig. 2g | C-fiber DREADD Gi PN male | 7 | -49 ± 3.63 | 78 ± 14 | Baseline: 0.9 ± 0.06 Post-CNO: 0.8 ± 0.06 | 0.054 | t=2.391, df=6 | Baseline: 102 ± 1.8 Post-CNO: 124 ± 3.7 | 0.0029 | t=4.848, df=6 |

**Supplementary Table 3. Behavior and calcium imaging.**

| Behavior - Gq female | | | | | | | |
|---|---|---|---|---|---|---|---|
| Fig. 3b | Von Frey (g) | Two-way RM ANOVA, Factor *Time* and *Treatment* (DREADD vs. Sham) | | | Time x Treatment P=0.0321 | F (6, 126) = 2.389 | |
| | | | | | Time P=0.0408 | F (4,228, 88,78) = 2.564 | |
| | | | | | Treatment P=0.1887 | F (1, 21) = 1.846 | |
| | | Baseline 1 | 31 ± 1.9 | 32 ± 1.9 | | | n.s. |
| | | Baseline 2 | 32 ± 2.1 | 35 ± 0.0 | | | n.s. |
| | | Baseline 3 | 29 ± 2.0 | 34 ± 1.4 | | | n.s. |
| | | 1-2 h | 20 ± 2.4 | 34 ± 1.4 | | | 0.0264 |
| | | 4-5 h | 25 ± 2.2 | 28 ± 3.2 | | | 0.9746 |
| | | 24 h | 27 ± 2.1 | 29 ± 2.1 | | | n.s. |
| | | 48 h | 26 ± 2.0 | 38 ± 2.1 | | | 0.8507 |
| **Behavior - Gi male** | | | | | | | |
| Fig. 3c | Von Frey (g) | Two-way RM ANOVA, Factor *Time* and *Treatment* (DREADD vs. Sham) | | | Time x Treatment P=0.0043 | F (6, 84) = 3.448 | |
| | | | | | Time P=0.0015 | F (6, 84) = 3.990 | |
| | | | | | Treatment P=0.0001 | F (1, 14) = 29.52 | |
| | | Baseline 1 | 24 ± 2.3 | 31 ± 1.9 | | | 0.3521 |
| | | Baseline 2 | 24 ± 2.3 | 28 ± 3 | | | 0.8631 |
| | | Baseline 3 | 23 ± 3.5 | 33 ± 1.7 | | | 0.0915 |
| | | 1-2 h | 7 ± 1.1 | 29 ± 2.6 | | | <0.0001 |
| | | 4-5 h | 16 ± 3.3 | 25 ± 3.5 | | | 0.1228 |
| | | 24 h | 23 ± 3.5 | 23 ± 2.9 | | | n.s. |
| | | 48 h | 20 ± 1.9 | 28 ± 2.5 | | | 0.2403 |
| **Behavior - Gi female** | | | | | | | |
| Fig. 3d | Von Frey (g) | Two-way RM ANOVA, Factor *Time* and *Treatment* (DREADD vs. Sham) | | | Time x Treatment P=0.0133 | F (6, 90) = 2.867 | |
| | | | | | Time P=0.0123 | F (4,240, 63,60) = 3.407 | |
| | | | | | Treatment P=0.0038 | F (1, 15) = 11.67 | |
| | | Baseline 1 | 22 ± 2.2 | 24 ± 2.0 | | | 0.9909 |
| | | Baseline 2 | 23 ± 2.0 | 26 ± 3.1 | | | 0.9891 |
| | | Baseline 3 | 21 ± 2.4 | 23 ± 2.3 | | | 0.9966 |
| | | 1-2 h | 8 ± 1.2 | 23 ± 2.3 | | | 0.0011 |
| | | 4-5 h | 16 ± 2.1 | 21 ± 1.5 | | | 0.3591 |
| | | 24 h | 22 ± 2.2 | 19 ± 2.7 | | | 0.9888 |
| | | 48 h | 21 ± 2.4 | 23 ± 2.3 | | | 0.9966 |

| Calcium Imaging | | | | | | |
|---|---|---|---|---|---|---|
| Figure | Group | Mean ± SEM | N (slices) | P-Value (2-way ANAOVA) | F | Post-hoc (adjusted P-Value) |
| Fig. 3h Events·min-1 | Sham Male | 0.356 ± 0.09 | 14 | Interaction: 0.4595<br>Treatment: <0.0001<br>Sex: 0.0020 | Interaction: F(2,94) = 0.7841<br>Treatment: F(2,94) = 13.31<br>Sex: F (1, 94) = 10.07 | Male:<br>Sham vs. Gq: 0.0001<br>Sham vs. Gi: 0.0005<br>Female:<br>Sham vs Gq: 0.0486<br>Sham vs Gi: 0.0278<br>Male - Female:<br>Sham: 0.8124<br>Gq: 0.0240<br>Gi: 0.1123 |
| | Gq Male | 1.215 ± 0.15 | 25 | | | |
| | Gi Male | 1.202 ± 0.25 | 16 | | | |
| | Sham Female | 0.179 ± 0.04 | 16 | | | |
| | Gq Female | 0.605 ± 0.11 | 15 | | | |
| | Gi Female | 0.620 ± 0.12 | 15 | | | |
| Fig. 3i Peak (ΔF/F) | Sham Male | 0.029 ± 0.01 | 14 | Interaction: 0.9134<br>Treatment: <0.0001<br>Sex: 0.3989 | Interaction: F(2,89) = 0.09065<br>Treatment: F(2,89) = 10.72<br>Sex: F (1, 89) = 0.7186 | Male:<br>Sham vs. Gq: 0.0058<br>Sham vs. Gi: 0.0056<br>Female:<br>Sham vs Gq: 0.0199<br>Sham vs Gi: 0.0206<br>Male - Female:<br>Sham: 0.9963<br>Gq: 0.9473<br>Gi: 0.8272 |
| | Gq Male | 0.098 ± 0.02 | 25 | | | |
| | Gi Male | 0.108 ± 0.03 | 16 | | | |
| | Sham Female | 0.025 ± 0.01 | 16 | | | |
| | Gq Female | 0.087 ± 0.02 | 15 | | | |
| | Gi Female | 0.067 ± 0.01 | 15 | | | |
| Fig. 3j Area (ΔF/F)ms | Sham Male | 0.338 ± 0.08 | 14 | Interaction: 0.3312<br>Treatment: <0.0001<br>Sex: 0.2338 | Interaction: F(2,89) = 1.119<br>Treatment: F(2,89) = 13.02<br>Sex: F (1, 89) = 1.437 | Male:<br>Sham vs. Gq: 0.0001<br>Sham vs. Gi: 0.9224<br>Female:<br>Sham vs Gq: 0.0189<br>Sham vs Gi: 0.4800<br>Male – Female:<br>Sham: 0.9518<br>Gq: 0.1528<br>Gi: 0.9954 |
| | Gq Male | 1.634 ± 0.30 | 25 | | | |
| | Gi Male | 0.463 ± 0.13 | 16 | | | |
| | Sham Female | 0.181 ± 0.08 | 16 | | | |
| | Gq Female | 1.042 ± 0.32 | 15 | | | |
| | Gi Female | 0.535 ± 0.16 | 15 | | | |

**Supplementary Table 4. Electrophysiological recordings in vivo (paired t-tests) and Perineuronal nets (one-way ANOVA).**

| Electrophysiology | | | | | |
|---|---|---|---|---|---|
| Figure | Treatment | Mean ± SEM (%) | N (animals) | P-Value (paired t-test) | t, df |
| Fig. 4b | Gonadectomized Male Gq-DREADD/CNO | Baseline: 99 ± 1<br>Post-CNO: 92 ± 14 | 7 | 0.6311 | t=0.5058, df=6 |
| Fig. 4c | Gonadectomized Male Gi-DREADD/CNO | Baseline: 101 ± 1<br>Post-CNO: 94 ± 15 | 7 | 0.6651 | t=0.4550, df=6 |
| Fig. 4d | Gonadectomized Female Gq-DREADD/CNO | Baseline: 100 ± 3<br>Post-CNO: 98 ± 6 | 8 | 0.6955 | t=0.4079, df=7 |
| Fig. 4e | Gonadectomized Female Gi-DREADD/CNO | Baseline: 101 ± 1<br>Post-CNO: 89 ± 12 | 7 | 0.3285 | t=1.064, df=6 |
| Fig. 4g | Testo Gonadectomized Female Gq-DREADD/CNO | Baseline: 98 ± 1<br>Post-CNO: 62 ± 7 | 11 | 0.0008 | t=4.725, df=10 |
| Fig. 4h | Testo Gonadectomized Female Gi-DREADD/CNO | Baseline: 100 ± 1<br>Post-CNO: 90 ± 15 | 11 | 0.5222 | t=0.6658, df=9 |

| Perineuronal nets | | | | | | | |
|---|---|---|---|---|---|---|---|
| Figure | Treatment | Mean ± SEM (%) | Animals | Slices | P-Value (one-way ANOVA) | F | Post-hoc (adjusted P-Value) |
| Fig. 4i | PNN mean intensity values (Aggrecan) whole SDH | Male: 30.03 ± 0.8921<br>Female + testo: 30.38 ± 1.393<br>Female + chABC: 30.7 ± 2.337<br>Female: 37.96 ± 2.085 | 4<br>3<br>2<br>2 | 16<br>16<br>13<br>10 | 0.0092 | F (3, 51) = 4.269 | Male vs. Female: 0.0133<br>Female vs. Female + testo: 0.0198<br>Male vs Female + chABC: 0.9999<br>Male vs Female + testo: >0.9999<br>Female vs Female + chABC: 0.0394<br>Female testo vs Female + chABC: >0.9999 |
| | PNN mean intensity values (WFA) whole SDH | Male: 31.82 ± 1.210<br>Female + testo: 35.87 ± 1.883<br>Female + chABC: 38.62 ± 2.813<br>Female: 51.77 ± 2.315 | 4<br>3<br>2<br>2 | 16<br>16<br>13<br>10 | <0.0001 | F (3, 51) = 15.55 | Male vs. Female: <0.0001<br>Female vs. Female + testo: <0.0001<br>Male vs Female + chABC: 0.1126<br>Male vs Female + testo: 0.8358<br>Female vs Female + chABC: 0.0008<br>Female testo vs Female + chABC: 0.1770 |

| Electrophysiology | | | | | |
|---|---|---|---|---|---|
| Figure | Treatment | Mean ± SEM (%) | N (animals) | P-Value (paired t-test) | t, df |
| Fig. 4l | ChABC Female Gq-DREADD/CNO | Baseline: 100 ± 1.48<br>Post-CNO: 42 ± 4.5 | 5 | <0.0001 | t=17.13, df=4 |
| Fig. 4m | ChABC Female Gi-DREADD/CNO | Baseline: 101 ± 1.22<br>Post-CNO: 97 ± 13.6 | 6 | 0.8370 | t=0.2167, df=5 |

**Supplementary Table 5: Membrane Potential, Cell Capacitance, Paired pulse ratio and evoked EPSCs of recorded cells.**

| Figure | Treatment | N (cells) | Resting Membrane Potential (mV) | Cell Capacitance (pF) | Paired Puls ratio (paired t-test) Mean ± SEM (%) | p-Value (paired t-test) | t, df | evoked EPSCs (paired t-test) Mean ± SEM (%) | p-Value (paired t-test) | t, df |
|---|---|---|---|---|---|---|---|---|---|---|
| Fig. 5a | C-fiber DREADD Gi BAPTA male | 9 | -45 ± 0.95 | 35 ± 4.8 | Baseline: 0.7 ± 0.07 Post-CNO: 0.7 ± 0.07 | 0.4556 | t=0.7840, df=8 | Baseline: 100 ± 0.8 Post-CNO: 96 ± 4.5 | 0.3729 | t=0.9437, df=8 |
| Fig. 5b | C-fiber DREADD Gi GDβS male | 7 | -45 ± 2.38 | 47 ± 13 | Baseline: 0.9 ± 0.06 Post-CNO: 0.8 ± 0.06 | 0.2248 | t=1.353, df=6 | Baseline: 96 ± 2.3 Post-CNO: 100 ± 9.4 | 0.7416 | t=0.3453, df=6 |
| Fig. 5c | C-fiber DREADD Gi DAP-5 male | 7 | -48 ± 1.59 | 46 ± 9.9 | Baseline: 0.7 ± 0.11 Post-CNO: 0.8 ± 0.1 | 0.3446 | t=1.026, df=6 | Baseline: 106 ± 2.1 Post-CNO: 82 ± 5 | 0.0023 | t=5.072, df=6 |

**Supplementary Table 6. Rostro-caudal spread.**

| Figure | Treatment | N animals | Rostro-caudal spread (mm) | N animals/slices | Area ($mm^2$) |
|---|---|---|---|---|---|
| Extended Data Fig. 1d | Male Gq | 3 | 3 ± 0.2 | 4/13 | 1.3 ± 0.1 |
| | Female Gq | 3 | 3.4 ± 1 | 4/16 | 1.3 ± 0.1 |
| | Male Gi | 3 | 3 ± 1 | 5/17 | 1.3 ± 0.1 |
| | Female Gi | 3 | 3 ± 0.3 | 4/16 | 1.4 ± 0.1 |

**Supplementary Table 7. Membrane Potential, Cell Capacitance, Paired pulse ratio and evoked EPSCs of recorded cells.**

| Figure | Treatment | N | Resting Membrane Potential (mV) | Cell Capacitance (pF) | Paired Puls ratio (paired t-test) Mean ± SEM (%) | p-Value (paired t-test) | t, df | evoked EPSCs (paired t-test) Mean ± SEM (%) | p-Value (paired t-test) | t, df |
|---|---|---|---|---|---|---|---|---|---|---|
| Extended Data Fig. 2a | Sham/ CNO Female | 5 | | | | | | Baseline: 99 ± 1 Post-CNO: 103 ± 8 | 0.7054 | t=0.4062, df=4 |
| Extended Data Fig. 2b | C-fiber DREADD Sham male | 7 | -46 ± 2.34 | 75 ± 17 | Baseline: 0.8 ± 0.1 Post-CNO: 0.8 ± 0.05 | 0.9737 | t=0.03438, df=6 | Baseline: 100 ± 1.4 Post-CNO: 93 ± 4.4 | 0.0905 | t=2.015, df=6 |
| Extended Data Fig. 2c | C-fiber DREADD Sham female | 7 | -47 ± 1.81 | 49 ± 6.3 | Baseline: 0.7 ± 0.14 Post-CNO: 0.6 ± 0.14 | 0.8229 | t=0.2338, df=6 | Baseline: 102 ± 1.2 Post-CNO: 102 ± 5.6 | 0.9686 | t=0.04102, df=6 |

### Supplementary Table 8. Cytokine Assay.

| Figure | Treatment (male vs female) | Mean ± SEM (%) | N (animals) | P-Value (paired T-Test) | t, df |
|---|---|---|---|---|---|
| Extended Data Fig. 5 - Gq | IL-13 | male: 0.78 ± 0.05<br>female: 1.29. ± 0.64 | 2<br>2 | 0.5650 | t=0.08108, df=1.012 |
| | IL-10 | male: 0.71 ± 0.18<br>female: 1.71 ± 0.46 | 2<br>2 | 0.2459 | t=2.018, df=1.294 |
| | IL-1a | male: 0.85 ± 0.66<br>female: 0.99 ± 0.57 | 2<br>2 | 0.8873 | t=0.1609, df=1.951 |
| | M-CSF | male: 1.11 ± 0.03<br>female: 1.1 ± 0.09 | 2<br>2 | 0.8943 | t=0.1630, df=1.155 |
| | IL-1b | male: 1.11 ± 0.06<br>female: 1.15 ± 0.34 | 2<br>2 | 0.9379 | t=0.09686, df=1.058 |
| | IFg | male: 0.84 ± 0.26<br>female: 1.04 ± 0.11 | 2<br>2 | 0.5654 | t=0.7430, df=1.345 |
| | MIP-3a | male: 0.96 ± 0.004<br>female: 1.02 ± 0.02 | 2<br>2 | 0.1988 | t=2.874, df=1.076 |
| | GmCsf | male: 0.99 ± 0.03<br>female: 0.93 ± 0.15 | 2<br>2 | 0.7814 | t=0.3504, df=1.097 |
| | IL-7 | male: 1.16 ± 0.22<br>female: 1.04 ± 0.08 | 2<br>2 | 0.6736 | t=0.5306, df=1.267 |
| | VEGF | male: 1.06 ± 0.03<br>female: 0.98 ± 0.17 | 2<br>2 | 0.7178 | t=0.4696, df=1.046 |
| | GRO/KC | male: 1.58 ± 0.52<br>female: 0.95 ± 0.07 | 2<br>2 | 0.4375 | t=1.197, df=1.038 |
| | IL-17A | male: 0.91 ± 0.16<br>female: 1.2 ± 0.11 | 2<br>2 | 0.2806 | t=1.543, df=1.736 |
| | IL-12p70 | male: 1.01 ± 0.01<br>female: 0.31 ± 0.06 | 2<br>2 | 0.1763 | t=3.434, df=1.021 |
| | IL-18 | male: 0.98 ± 0.004<br>female: 1.02 ± 0.08 | 2<br>2 | 0.6805 | t=0.5480, df=1.005 |
| | IL-2 | male: 1.01 ± 0.12<br>female: 1.254 ± 0.27 | 2<br>2 | 0.5235 | t=0.8322, df=1.396 |
| | TNF-a | male: 1.22 ± 0.22<br>female: 0.79 ± 0.38 | 2<br>2 | 0.4456 | t=0.9961, df=1.602 |
| | MCP-1 | male: 1.24 ± 0.15<br>female: 1.06 ± 0.15 | 2<br>2 | 0.4657 | t=0.8942, df=1.996 |
| | IL-6 | male: 1.12 ± 0.12<br>female: 0.93 ± 0.01 | 2<br>2 | 0.3641 | t=1.544, df=1.010 |
| | MIP1a | male: 2.47 ± 1.31<br>female: 1.18 ± 0.19 | 2<br>2 | 0.5020 | t=0.9769, df=1.043 |
| Extended Data Fig. 5 - Gi | IL-13 | male: 1.08 ± 0.04<br>female: 1.07 ± 0.12 | 2<br>2 | 0.9778 | t=0.03380, df=1.197 |
| | IL-10 | male: 1.24 ± 0.41<br>female: 1.53 ± 0.12 | 2<br>2 | 0.6093 | t=0.6713, df=1.178 |
| | IL-1a | male: 1.77 ± 0.2<br>female: 1.44 ± 0.55 | 2<br>2 | 0.6539 | t=0.5697, df=1.255 |
| | M-CSF | male: 1.14 ± 0.12<br>female: 1.01 ± 0.1 | 2<br>2 | 0.4958 | t=0.8273, df=1.981 |
| | IL-1b | male: 1.505 ± 0.01<br>female: 1.66 ± 0.34 | 2<br>2 | 0.7292 | t=0.4529, df=1.001 |

| | | | | | |
|---|---|---|---|---|---|
| | IFg | male: 1.17 ± 0.38<br>female: 1.02 ± 0.29 | 2<br>2 | 0.7866 | t=0.3114, df=1.882 |
| | MIP-3a | male: 1.19 ± 0.03<br>female: 1.02 ± 0.05 | 2<br>2 | 0.1332 | t=2.799, df=1.642 |
| | GmCsf | male: 1.07 ± 0.12<br>female: 1.11 ± 0.02 | 2<br>2 | 0.7866 | t=0.3455, df=1.036 |
| | IL-7 | male: 1.21 ± 0.06<br>female: 1.23 ± 0.08 | 2<br>2 | 0.8630 | t=0.1974, df=1.856 |
| | VEGF | male: 1.19 ± 0.12<br>female: 1.17 ± 0.01 | 2<br>2 | 0.8786 | t=0.1929, df=1.004 |
| | GRO/KC | male: 1.76 ± 0.31<br>female: 1.19 ± 0.004 | 2<br>2 | 0.3227 | t=1.801, df=1.000 |
| | IL-17A | male: 1.15 ± 0.28<br>female: 0.98 ± 0.29 | 2<br>2 | 0.7131 | t=0.4236, df=1.995 |
| | IL-12p70 | male: 1.35 ± 0.17<br>female: 1.31 ± 0.19 | 2<br>2 | 0.8980 | t=0.1452, df=1.975 |
| | IL-18 | male: 1.03 ± 0.05<br>female: 1.02 ± 0.04 | 2<br>2 | 0.8646 | t=0.1937, df=1.956 |
| | IL-2 | male: 1.03 ± 0.25<br>female: 0.68 ± 0.03 | 2<br>2 | 0.3926 | t=1.391, df=1.026 |
| | TNF-a | male: 2.24 ± 0.55<br>female: 1.56 ± 0.13 | 2<br>2 | 0.4186 | t=1.229, df=1.114 |
| | MCP-1 | male: 1.25 ± 0.19<br>female: 1.06 ± 0.07 | 2<br>2 | 0.4931 | t=0.9408, df=1.246 |
| | IL-6 | male: 1.12 ± 0.02<br>female: 1.03 ± 0.12 | 2<br>2 | 05843 | t=0.7580, df=1.028 |
| | MIP1a | male: 2.71 ± 0.6<br>female: 1.33 ± 0.31 | 2<br>2 | 0.2174 | t=2.065, df=1.484 |

**Supplementary Table 9. Androgen receptor analysis (unpaired t-tests), electrophysiology, and testosterone measurements**

| Androgen receptor analysis | | | | | | |
|---|---|---|---|---|---|---|
| Figure | Treatment | Mean ± SEM (%) | Animals | Slices | P-Value (paired t-test) | t, df |
| Extended Data Fig. 6b | % $NeuN^+/AR^+$ | Male: 84.37 ± 2.035<br>Female: 73.04 ± 5.075 | 2<br>2 | 6<br>6 | 0.0652 | t=2.071, df=10 |
| Extended Data Fig. 6c | % $GFAP^+$-$S100b^+/AR^+$ | Male: 84.30 ± 1.654<br>Female: 77.35 ± 4.413 | 2<br>2 | 6<br>6 | 0.1714 | t=1.474, df=10 |
| Extended Data Fig. 6d | $AR^+$ cells/$\mu m^3$ | Male: 180.0 ± 14.07<br>Female: 154.3 ± 10.77 | 2<br>2 | 6<br>6 | 0.1781 | t=1.449, df=10 |

| Electrophysiology | | | | | |
|---|---|---|---|---|---|
| Figure | Treatment | Mean ± SEM (%) | N (animals) | P-Value (paired t-test) | t, df |
| Extended Data Fig. 6e | Non-OVX female + testo Gq-DREADD/CNO | Baseline: 98 ± 1.45<br>Post-CNO: 47 ± 13.3 | 5 | 0.0144 | t=4.136, df=4 |

| Testosterone measurement | | | |
|---|---|---|---|
| Figure | Treatment | Mean ± SEM (ng/µl) | Animals |
| Extended Data Fig. 6f | Testosterone analysis | Male: 2.350 ± 0.5914<br>Female: 6.404 ± 0.3604 | 4<br>24 |

**Supplementary Table 10. Antibodies and reagents used in present study.**

| Antibody | Dilution | Species | Cat. Nr. | Company |
|---|---|---|---|---|
| Aggrecan | 1:75 | Mouse | MA316888 | Invitrogen |
| Androgen Receptor | 1:150 | Rabbit | PA1-110 | Invitrogen |
| GFAP | 1:1000 | Chicken | PA1-10004 | Thermo Fisher Scientific |
| GFAP | 1:500 | Rabbit | 12389 | Cell Signaling |
| NeuN | 1:600 | Mouse | MAB377 | Milipore |
| NeuN | 1:1000 | Rabbit | 12943S | Cell Signalling |
| NK1 | 1:1000 | Guinea pig | AB15810 | Millipore |
| S100beta | 1:500 | Guinea pig | 287004 | Synaptic Systems |
| Wisteria Floribunda Lectin | 1:1000 | biotin | B-1355-2 | Vector Laboratories |
| Alexa Flour 488 | 1:500 | Goat α Guinea pig | A11073 | Thermo Fisher Scientific |
| Alexa Flour 488 | 1:500 | Goat α Chicken | A11039 | Thermo Fisher Scientific |
| Alexa Flour 488 | 1:250 | Streptavidin | S32354 | Thermo Fisher Scientific |
| Alexa Flour 488 | 1:1000 | Goat α Rabbit | A11008 | Thermo Fisher Scientific |
| Alexa Flour 546 | 1:500 | Goat α Mouse | A11003 | Thermo Fisher Scientific |
| Alexa Flour 647 | 1:250 | Goat α Guinea pig | A21450 | Thermo Fisher Scientific |
| Alexa Flour 647 | 1:500 | Goat α Rabbit | A21245 | Thermo Fisher Scientific |
| Alexa Flour 647 | 1:1000 | Goat α Mouse | A21235 | Invitrogen |
| BrilliantViolet 421 | 1:100 | Donkey α Rabbit | 406410 | BioLegend |
| BrilliantViolet 421 | 1:1000 | Streptavidin | 405226 | BioLegend |
